# All-optical electrophysiology reveals excitation, inhibition, and neuromodulation in cortical layer 1

**DOI:** 10.1101/614172

**Authors:** Linlin Z. Fan, Simon Kheifets, Urs L. Böhm, Kiryl D. Piatkevich, Hao Wu, Vicente Parot, Michael E. Xie, Edward S. Boyden, Anne E. Takesian, Adam E. Cohen

## Abstract

The stability of neural dynamics arises through a tight coupling of excitatory (E) and inhibitory (I) signals. Genetically encoded voltage indicators (GEVIs) can report both spikes and subthreshold dynamics *in vivo*, but voltage only reveals the combined effects of E and I synaptic inputs, not their separate contributions individually. Here we combine optical recording of membrane voltage with simultaneous optogenetic manipulation to probe E and I individually in barrel cortex Layer 1 (L1) neurons in awake mice. Our studies reveal how the L1 microcircuit integrates thalamocortical excitation, lateral inhibition and top-down neuromodulatory inputs. We develop a simple computational model of the L1 microcircuit which captures the main features of our data. Together, these results suggest a model for computation in L1 interneurons consistent with their hypothesized role in attentional gating of the underlying cortex. Our results demonstrate that all-optical electrophysiology can reveal basic principles of neural circuit function *in vivo*.

**One Sentence Summary:** All-optical electrophysiology revealed the function in awake mice of an inhibitory microcircuit in barrel cortex Layer 1.

## Main Text

The brain receives myriad sensory inputs. It must distinguish the relevant from the irrelevant. An input can merit attention either through its intrinsic properties (novelty, salience) or through learned associations. The sparse interneurons of neocortical Layer 1 (L1) have been hypothesized as a hub for integrating these factors and modulating the underlying cortex to control attention (*1–3*). L1 interneurons receive direct thalamic (*2, 4, 5*), cortico-cortical (*6, 7*), and neuromodulatory (cholinergic (*2, 3, 8*) and serotonergic (*8, 9*)) inputs, the last mediated by fast ionotropic receptors. Activation of L1 interneurons exerts powerful control of underlying cortex by inhibiting deeper-lying interneurons and thereby dis-inhibiting pyramidal neurons (*1–3, 6*). L1 interneurons also directly inhibit pyramidal neuron apical dendrites (*2, 10, 11*). Despite the suggestive anatomy and influence on underlying cortex, little is known about information processing within L1, especially in awake animals.

A core principle of neuronal network dynamics is maintenance of balance between excitation (E) and inhibition (I). For instance, during sensory processing in cortical layer 4 (L4) excitation driven by thalamic inputs is countered by feed-forward inhibition from parvalbumin (PV) interneurons (*12, 13*). L1 interneurons receive inhibitory inputs both from other L1 interneurons (*5, 14, 15*) and from deeper Martinotti cells (*11*). It is not known how these inputs influence L1 activity *in vivo*.

Electrophysiological studies in L1 *in vivo* have been challenging due to the sparseness of neuronal cell bodies. While a few whole-cell patch clamp recordings have been performed in anesthetized rats (*1, 4*), technical difficulties have prevented similar acquisitions in awake animals. Due to their multimodal, temporally precise inputs (*16*) and temporally precise outputs, one would like to measure the sub-threshold dynamics and spike timing of L1 neurons with high precision in voltage and time. Recent advances in genetically encoded voltage indicators (GEVIs) enabled voltage imaging with single-neuron, single-spike resolution in hippocampus (*17–20*) and in superficial cortex (*18, 19*) *in vivo*, opening the possibility for optical explorations of L1 circuit function *in vivo*.

Voltage alone does not distinguish the relative contributions of E and I synaptic inputs, yet this distinction is critical for understanding circuit mechanisms. A commonly used patch clamp technique is to inject current to achieve different levels of baseline depolarization and thereby to shift the relative driving force of E vs. I post-synaptic currents, revealing their distinct contributions (*21*). We previously paired near infrared GEVIs based on Archaerhodopsin 3 (Arch) with channelrhodopsin stimulation for optical measurements of excitability *in vivo* (Optopatch) (*17, 22*) and of synaptic transmission in primary culture and acute slices (*23*). Here we show that optogenetic depolarization of a postsynaptic neuron during sensory processing *in vivo* can unmask otherwise hidden inhibitory inputs.

We combine a novel holographic structured illumination imaging system, an Archaerhodospsin-derived GEVI optimized for crosstalk-free *in vivo* Optopatch, and patterned optogenetic stimulation to study the role of excitatory, inhibitory and neuromodulatory inputs on the function of the cortical L1 microcircuit during sensory processing in awake mice. A simple computational model that incorporates the known morphology, electrophysiology, and connectivity of L1 interneurons captures the main features of our data and suggests an intuitive picture for novelty detection in L1.

## Results

### *In vivo* Optopatch with holographic patterned illumination

Archon1 is an Arch-derived GEVI with improved trafficking and brightness (*24*). A soma-localized variant, SomArchon, enabled voltage imaging *in vivo* with good signal-to-noise ratio (*20*). We made a Cre-dependent bicistronic construct for co-expression of SomArchon and a blue light-activated soma-localized channelrhodopsin, CheRiff (*25*). We call this combined construct Optopatch4.

Voltage signals in tissue arise solely from the neuronal membrane. Illumination that enters the tissue but misses the membrane of interest contributes to background fluorescence and heating, but not to signal. In epifluorescence images of membrane-labeled neurons, the soma perimeter appears brighter than the center, a geometrical projection effect from viewing membranes edge-on. We thus reasoned that incident photons would most efficiently produce signal if targeted to the soma perimeter. Confocal-like excitation combined with spatially filtered emission would also minimize optical crosstalk from out-of-focus cells. We built a holographic structured illumination system, similar to Ref. (*26*), to achieve this precisely targeted illumination with red (*λ* = 635 nm) light for excitation of SomArchon (Fig. 1A, Fig. S1, Table S3, Methods). SomArchon fluorescence from all holographically targeted spots was recorded simultaneously on a scientific CMOS camera. Spatial filters were applied digitally in post-processing to separate signal from background (Methods). A digital micromirror device (DMD) patterned blue illumination for targeted optogenetic stimulation (Fig. 1A, Fig. S1, Table S3).

**Figure 1.**
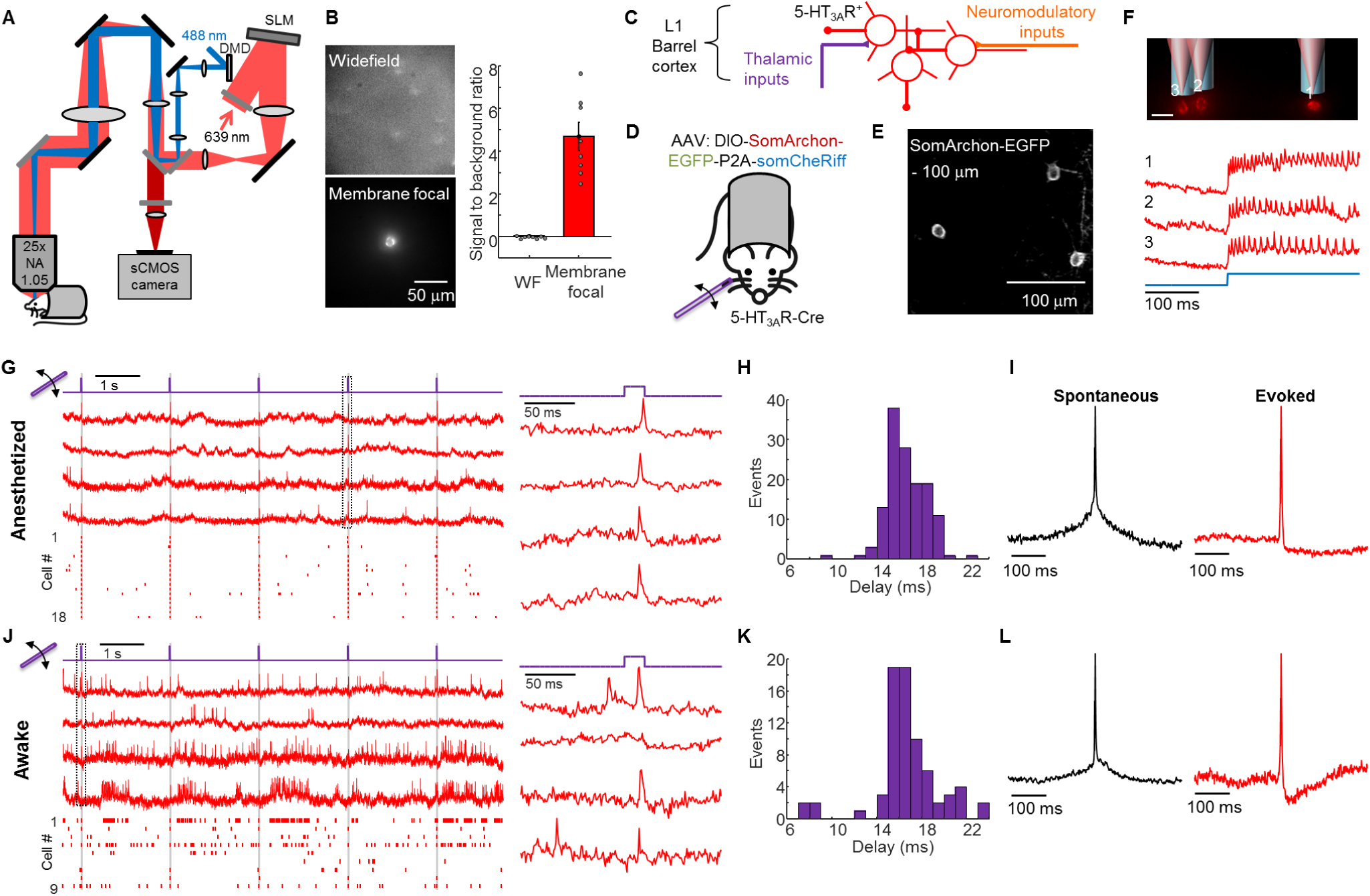
All-optical electrophysiology in L1 neurons *in vivo*. (**A**) Optical system for holographic structured illumination voltage imaging (red light) and micromirror-patterned optogenetic stimulation (blue light). Details in **Methods**. (**B**) Comparison of widefield epifluorescence and membrane-targeted holographic illumination of the same field of view containing a SomArchon-expressing L1 neuron. Scale bar 50 µm. Right: quantification of the signal-to-background ratio for the two imaging modalities. (**C)** 5-HT_3A_R-positive interneurons in L1 of the barrel cortex receive sensory inputs from the thalamus, neuromodulatory inputs from higher brain regions, and lateral inhibition from other L1 interneurons. (**D**) Virus encoding a cre-dependent construct for co-expression of a voltage indicator (SomArchon-EGFP) and an optogenetic actuator (somCheRiff) were injected into barrel cortex of 5-HT_3A_R-Cre mice. A glass capillary delivered small stimuli to an individual whisker. (**E**) Two-photon microscopy image of GFP fluorescence from SomArchon-EGFP in barrel cortex L1 showing good trafficking and soma localization. Scale bar: 100 µm, image depth 100 µm below dura. (**F**) Combination of patterned optogenetic stimulation (blue) and holographic illumination for voltage imaging. Scale bar 20 µm. Bottom: fluorescence traces from the three indicated cells in response to a step in blue illumination in an anesthetized mouse. (**G**) Fluorescence transients in single L1 interneurons evoked by whisker stimuli (20 ms deflections at 0.5 Hz) in anesthetized mice. Left, top: examples fluorescence traces recorded at 1 kHz frame rate. Traces have been corrected for photobleaching but not otherwise filtered. Bottom: raster plot showing spikes from *n* = 18 neurons. Right: Fluorescence waveforms from the boxed region at left. (**H**) Distribution of delays between stimulus onset and peak of evoked spike. (**I**) Spike-triggered average waveform of spontaneous (left, *n* = 17 neurons) and whisker stimulus-evoked (right, *n* = 24 neurons) action potentials. A spike was classified as ‘evoked’ if it occurred within 30 ms of stimulus onset. (**J-L**) Same as **G-I** but in an awake mouse. Spontaneous: *n* = 22 neurons. Evoked: *n* = 21 neurons. Data in **G-L** recorded from 3 mice.

We characterized the performance of the system by imaging SomArchon-expressing neurons *in vivo* in the cortex. Under wide-field red illumination, the cells were not visible due to high background from scattered light (Fig. 1B). Illumination targeted to the somas revealed individual cells (Fig. 1B, Fig. S2). Holographic membrane-targeted illumination provided substantially better optical sectioning and signal-to-background ratio than did soma-wide illumination (Fig. S2). The combination of SomArchon and the holographic optical system enabled recording of spontaneous action potentials with SNR 12 ± 4 (mean ± s.d., *n* = 16 cells) at depths between 20 and 150 µm and SNR 6.7 at a depth of ∼200 µm at a 1 kHz frame rate in awake head-fixed mice (Fig. S3, Methods).

To target expression to L1, we expressed Optopatch4 in 5HT_3A_R-Cre mice (Fig. 1C, D). This line drives expression predominantly in supragranular layers, including in ∼90% of L1 interneurons (*2, 9*). Two-photon fluorescence images of an appended eGFP tag showed membrane-localized and somatically restricted expression in L1 (Fig. 1E). In acute slices, targeted optogenetic stimuli evoked characteristic firing patterns in L1 neurons, including previously reported bursting adapting and late-spiking non-adapting phenotypes (*14*) (Fig. S4), though not all neurons had a clear electrophysiological classification.

In head-fixed mice, targeted optogenetic stimuli evoked spikes which were clearly resolved via holographically targeted voltage imaging in recordings acquired at a 1 kHz frame-rate (Fig. 1F). We measured excitability and firing properties of L1 neurons (depth < 150 µm) in mice anesthetized with isoflurane, and then later re-measured the same neurons in awake mice (Fig. S5). While awake mice tended to show higher excitability and more variable subthreshold dynamics, the core firing properties (e.g. bursting, adaptation) were qualitatively preserved within each cell between the two brain states (Fig. S5). Consistent with prior reports of *in vivo* patch clamp measurements in L1 interneurons (*27, 28*), the firing patterns did not clearly resolve into distinct sub-classes. We therefore treated all measured neurons as a single population.

### Voltage imaging of whisker stimulus-triggered activity in L1 neurons

Barrel fields corresponding to individual whiskers (B2, C2, D2) were identified by intrinsic imaging (Fig. S6, Methods). We then used voltage imaging to characterize the sensory-evoked responses in L1 interneurons. In both anesthetized and awake mice, brief stimuli to individual whiskers (∼1 mm deflection, ∼8 mm from the base, 20 ms duration, repeated at 0.5 Hz, Methods) elicited excitatory post-synaptic potentials (EPSPs) and often spikes in L1 neurons in the corresponding barrel fields (Fig. 1G, H). The delay from stimulus onset to spike peak was 16 ± 2 ms (mean ± s.d., *n* = 135 events, 24 neurons, 3 mice) in anesthetized mice and 16 ± 3 ms in awake mice (mean ± s.d., *n* = 73 events, 21 neurons, 3 mice, Fig. 1J, K). Similar delay and jitter were previously reported in L4 pyramidal neurons and fast-spiking neurons, both of which receive direct thalamic inputs (*12*).

In a comparison between spontaneous and whisker-evoked spikes, we observed striking differences in the mean subthreshold dynamics calculated via a spike-triggered average (STA, Fig. 1I, L). Spontaneous spikes rode atop a baseline depolarization that both preceded and followed the spike, whereas whisker-evoked spikes arose abruptly and were followed by a period of hyperpolarization (Fig. 1I, L). Stimulus-triggered average waveforms of whisker deflection trials that did not induce spikes also showed a depolarization followed by a hyperpolarization (Fig. S7). Together, these results implied that the subthreshold dynamics preceding and following spikes reflected distinct network states for spontaneous and sensory-evoked activity, rather than purely cell-autonomous effects of voltage-gated ion channels.

STA waveforms also differed between anesthetized and awake animals (Fig. 1I, L). For spontaneous spikes, the subthreshold depolarization was larger under anesthesia than wakefulness (anesthetized: 22 ± 2% of spike height, *n* = 17 neurons, 3 mice vs. awake: 10 ± 2% of spike height, *n* = 22 neurons, 3 mice, *p* = 3 × 10^−4^, two-tailed *t*-test, mean ± s.e.m.). For whisker-evoked spikes, the after-spike hyperpolarization was smaller but longer lasting under anesthesia than under wakefulness than (anesthetized: 11 ± 2% spike height, *n* = 24 neurons, 3 mice vs. awake: 17 ± 1% of spike height, *n* = 21 neurons, 3 mice, *p* = 0.02, two-tailed *t*-test; anesthetized: 254 ± 15 ms recovery time vs. awake: 127 ± 14 ms, *p* = 1 × 10^−7^, two-tailed *t*-test, all mean ± s.e.m). These observations are consistent with a more depolarized resting potential under wakefulness, consistent with prior reports (*29*).

### Optical dissection of excitation and inhibition during sensory processing

Rapid inhibition is mediated by GABAA receptors, ligand-gated chloride channels with a reversal potential of ∼-70 mV. L1 interneurons in anesthetized rats have been reported to rest at −65 to −70 mV (*4*), suggesting that inhibitory inputs should have only small effects on membrane potential at rest. Borrowing from well-established patch clamp protocols (*21*), we reasoned that optogenetic depolarization would increase the driving force for inward chloride current, and thereby amplify the impact of GABAA receptor activation on the inhibitory postsynaptic potential (IPSP) (Fig. 2A,B).

**Figure 2.**
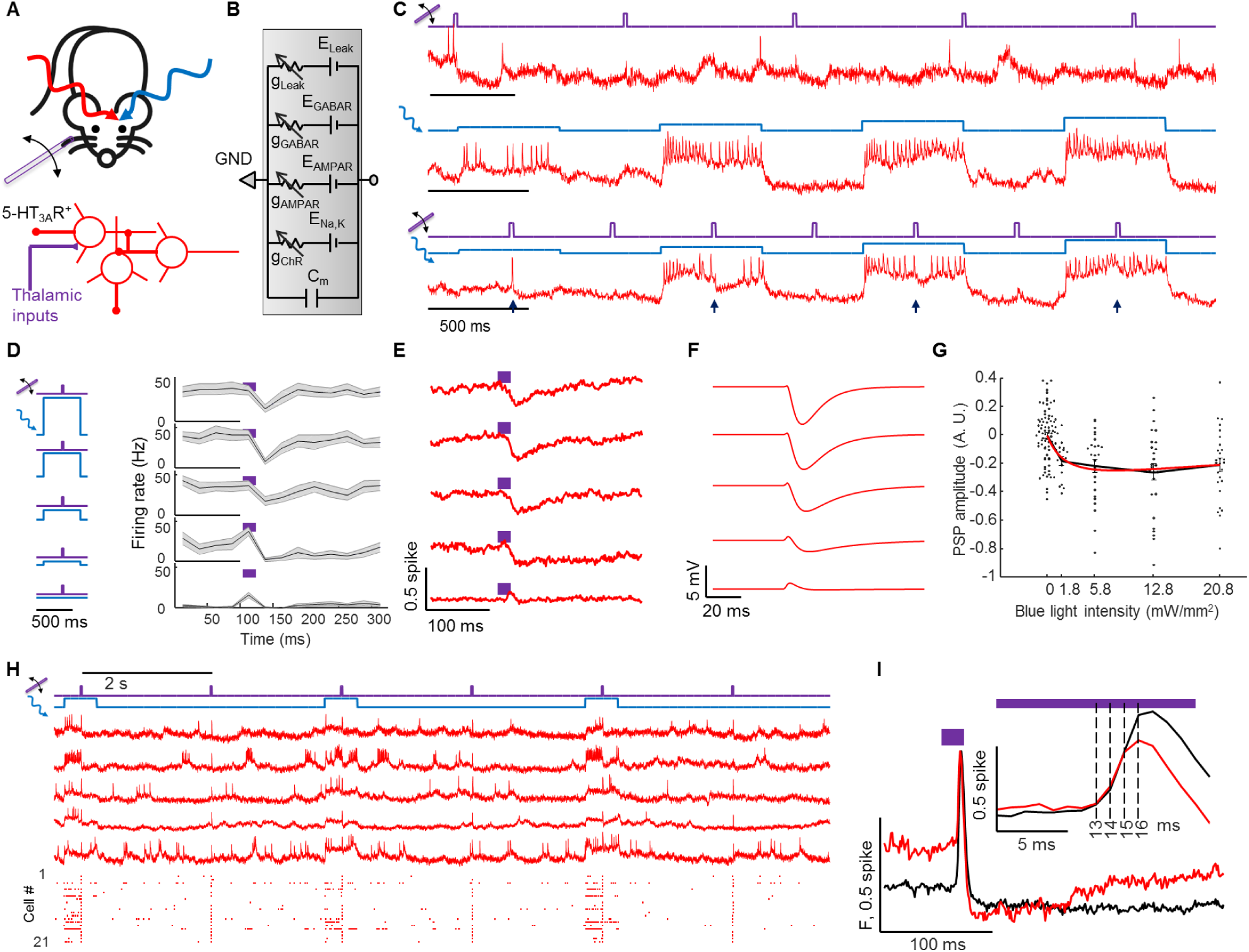
Optical dissection of excitation and inhibition in L1 interneurons in awake mice. (**A**) Whisker stimuli and single cell-targeted optogenetic stimuli were paired in 5HT_3A_R-Cre mice expressing Optopatch4. (**B**) Conductance-based model of membrane potential. This simple model only contains passive conductances, with gating by light (Channelrhodopsin, ChR), glutamate (AMPAR), and GABA (GABAR). A leak conductance sets the resting potential of the cell in the absence of optogenetic or synaptic inputs. (**C**) Three recordings from a single neuron showing response to (top) whisker stimulus, (middle) optogenetic stimulus, and (bottom) simultaneous optogenetic and whisker stimuli. Arrows show whisker stimulus-evoked inhibition. (**D**) Mean spike rate evoked by whisker stimuli atop different levels of optogenetic stimulus. In the absence of optogenetic stimulation, whisker stimuli evoked precisely timed single spikes. In the presence of optogenetic stimulation, whisker stimuli suppressed spiking. The suppression decreased in amplitude and duration as the strength of the optogenetic stimulus increased. Shading represents s.e.m. from *n* = 27 neurons, 4 mice. (**E**) Mean whisker stimulus-evoked subthreshold waveforms at different levels of optogenetic drive. Spikes were digitally removed prior to averaging (Methods). (**F**) Simulated membrane voltage waveforms under different levels of optogenetic drive, using the model shown in (B). Excitation was assumed to lead inhibition by 2 ms. Details in Methods. (**G**) Comparison of PSP amplitude as a function of optogenetic stimulus strength with numerical simulation from a simple conductance-based model. (**H**) Repetitive measurements of whisker stimulus-evoked responses in anesthetized mice, with and without baseline optogenetic stimulation. Top: example recordings. Bottom: spike raster from *n* = 21 neurons, 3 mice. (**I**) Mean fluorescence responses to whisker stimulus without (black) and with (red) baseline optogenetic stimulation. Traces have been aligned to their peak. Inset: delay between onset of excitation (between 13 and 14 ms) and onset of inhibition (between 15 and 16 ms). To facilitate comparison, traces were vertically offset to align baseline values.

In both awake and anesthetized mice, whisker stimuli in the absence of optogenetic stimulation evoked clear spikes or EPSPs in L1 interneurons, as in prior experiments (Fig. 2C, S8). Optogenetic stimuli targeted to the same cells one at a time (500 ms duration, 1.8 to 21 mW/mm^2^, repeated at 1 Hz) reliably evoked stimulus intensity-dependent spiking. Remarkably, whisker stimuli applied during targeted single-cell optogenetic stimulation led to suppressed spiking, and hyperpolarization (Fig. 2C, D, E, S8). We quantified the sensory-evoked subthreshold waveforms by digitally removing spikes (Methods) and calculating a stimulus-triggered average at different optogenetic stimulus strengths (Fig. 2E, S8). In both awake and anesthetized brain states, whisker stimuli had opposite effects in the absence vs. presence of baseline optogenetic stimulation, illustrating dramatic non-additivity of sensory and optogenetic inputs to the same neuron.

A simple biophysical model containing passive leak, channelrhodopsin, AMPA receptor and GABAA receptor conductances captured the main features of our data (Fig. 2F, G, Methods). We assumed a transient excitatory synaptic input followed shortly by a transient inhibitory input. The model confirmed that optogenetic depolarization increased the driving force for chloride, revealing the presence of otherwise hidden sensory-evoked inhibitory inputs. Depolarization via endogenous currents, as occurred in the transition from anesthesia to wakefulness, also amplified the impact of transient inhibition, explaining the difference in whisker-evoked subthreshold waveforms in Figs. 1I, L.

Despite lacking many details (e.g. active conductances), the biophysical model captured several subtle aspects of the subthreshold dynamics. In the anesthetized state, as the strength of the optogenetic drive increased, the sensory-evoked IPSP amplitude first increased—as explained above—but then decreased (IPSP amplitude 29 ± 5% of spike height at low optogenetic drive (5.8 mW/cm^2^) vs. 16 ± 4% of spike height at high optogenetic drive (21 mW/cm^2^), *n* = 15 neurons, 3 mice, *p* = 0.001, two-sided paired-sample *t*-test, mean ± s.e.m.). A similar, but not statistically significant, trend occurred in the awake state (IPSP amplitude 28 ± 4% of spike height at low optogenetic drive vs. 23 ± 4% of spike height at high optogenetic drive, *n* = 27 neurons, 4 mice, *p* = 0.19, two-sided paired-sample *t*-test). The model revealed that these decreases were due to shunting of the membrane potential toward CheRiff reversal potential at high CheRiff conductance (∼0 mV, Supplementary Text). The IPSP duration also became shorter under strong optogenetic drive in both awake and anesthetized brain states. The model ascribed this effect to a decreased membrane RC time-constant due to the high CheRiff conductance. Our simple model thus connected the complex context-dependent whisker-evoked responses in L1 interneurons to basic membrane biophysics.

### Temporal dissection of excitation and inhibition

We next asked about the relative timing of excitatory and inhibitory inputs. We delivered whisker stimuli alternately with and without baseline weak optogenetic stimulation targeted to single neurons (5.8 mW/mm^2^, Fig. 2H). We anticipated that the whisker-evoked responses in these two conditions would initially coincide and then would diverge upon arrival of the inhibitory inputs. We compared stimulus-triggered average waveforms of trials that evoked spikes (Fig. 2I). The shape of the waveforms overlapped for the first 2 ms after onset of whisker-evoked depolarization. Thereafter, the waveform in the presence of optogenetic stimulation fell below the waveform in the absence, signaling the onset of inhibition (Fig. 2I, inset). This finding implies a ∼2 ms delay between onset of excitation and inhibition, suggesting at most a difference of one synapse in the respective paths (*12*). This result does not rule out the possibility that slower inhibitory signals (e.g. from GABAB receptors or polysynaptic mechanisms) also contributed to inhibition at later times.

### Lateral inhibition

We sought to identify the source of the sensory-evoked inhibition. Patch clamp measurements in acute slices have identified inhibitory connections between L1 interneurons (*5, 14, 15*). Since our whisker stimuli often evoked spikes in arbitrarily selected L1 interneurons, and inhibition lagged excitation by only ∼2 ms, we hypothesized that the rapid whisker stimulus-evoked inhibition was due to lateral connections within the L1 population.

To test this hypothesis, we performed an all-optical circuit-mapping experiment *in vivo* using patterned optogenetic stimulation (Fig. 3A). We expressed Optopatch4 in 5-HT_3A_R-Cre mice and targeted voltage imaging to 1-3 L1 interneurons in the center of the field of view. We then defined two optogenetic stimulus patterns. The first pattern comprised small disks targeted individually to the central neurons. These disks were stimulated with long pulses of blue light (500 ms, 25 mW/mm^2^), with the goal to depolarize the targeted cells and to increase the driving force for inhibitory currents. The second pattern comprised an annulus (inner diameter ∼200 µm, outer diameter of ∼400 µm, Fig. 3B, C, Methods), surrounding the central neurons. Midway through the stimulation of the central neurons, a brief flash (20 ms, 25 mW/mm^2^) was applied to the neurons in the surrounding annulus to evoke synchronized spiking of the surrounding cells.

**Figure 3.**
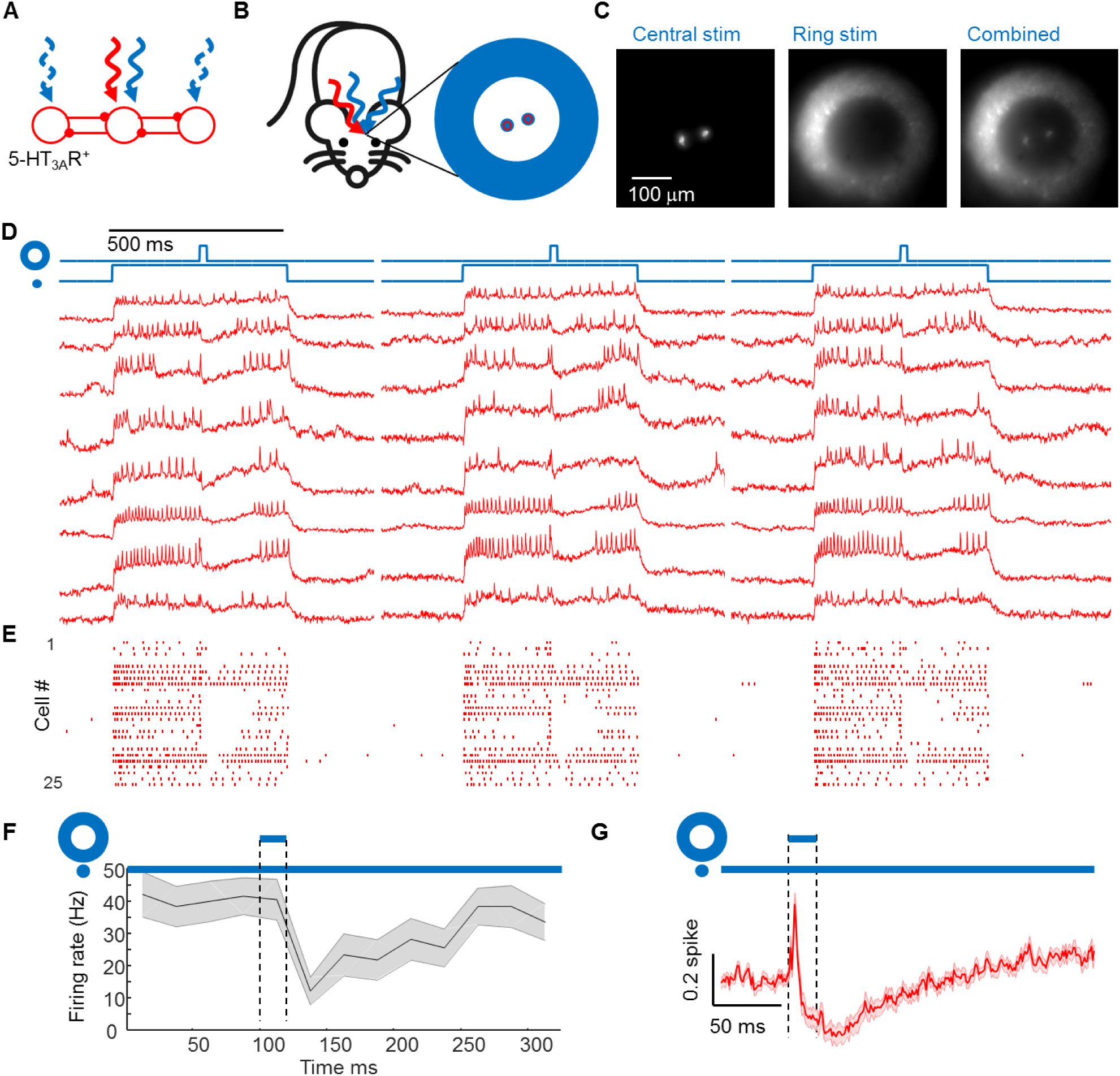
Center/surround optogenetic stimulation reveals lateral inhibition in L1. (**A**) Simple model of L1 circuit with lateral inhibition. Tonic optogenetic stimulation depolarizes the central neuron, increasing the driving force for inhibitory currents. Pulsed optogenetic stimulation of the surrounding neurons evokes lateral inhibition, revealed by voltage imaging (red) in the central neuron. (**B**) Experiment to probe lateral inhibition in L1. 5-HT_3A_R-Cre mice expressed Optopatch4 in barrel cortex. Optogenetic stimuli were delivered separately to central and surrounding neurons. Voltage imaging was performed only in central neurons. Experiments were performed in anesthetized mice. (**C**) Epifluorescence images showing the illumination patterns *in vivo*. Scale bar 100 µm. (**D**) Fluorescence waveforms from the central neurons under center/surround optogenetic stimulation. Central stimulation depolarized the targeted neurons and evoked spiking. Surround stimulation hyperpolarized the targeted neurons and suppressed spiking. (**E**) Spike raster showing responses from *n* = 25 neurons, 3 mice. (**F**) Mean spike rate during central stimulation, before and after surround stimulation. Surround stimulation caused spike rate to drop from 40.5 ± 6.3 Hz to 12.3 ± 4.1 Hz, *n* = 25 neurons, 3 mice (*p* = 4×10^−4^, two-sided paired-sample t-test). Shading represents s.e.m. (**G**) Mean subthreshold voltage during central stimulation, before and after surround stimulation. Surround stimulation caused inhibition in the central neuron. The initial spike in membrane voltage in the central neuron was due to scattered light from the surround which drove direct CheRiff activation. Shading represents s.e.m..

Optogenetic stimulation of the central neurons evoked robust spiking (spike rate 41 ± 6 Hz, *n* = 25 neurons, 3 mice, mean ± s.e.m.). Stimulation of the surrounding neurons transiently suppressed this spiking (spike rate 12 ± 4 Hz in the 25 ms following the annular flash, *p* = 4 × 10^−4^, two-sided paired-sample *t*-test, Fig. 3D, E, F). The mean fluorescence waveform following the annular flash showed robust hyperpolarization of the central neurons (27 ± 3% of spike height, Fig. 3G, S9). (Control experiments without the central optogenetic stimulus revealed that the initial depolarization after the annular flash was an artifact from light scatter, Fig. S9). The spike patterns and subthreshold hyperpolarization dynamics in these experiments closely resembled the corresponding data for a sensory stimulus (Fig. 2E, F, I). These results are consistent with the model that sensory stimulation elicits rapid activation of L1 neurons followed by rapid lateral inhibition within the L1 microcircuit.

### Neuromodulation

Finally, we explored the role of neuromodulatory activity on L1 dynamics (Fig. 4A). A mild air puff to the face has been shown to activate cholinergic neurons in basal forebrain (*16*), and these neurons are known to innervate cortical L1 (*3, 30*). We imaged L1 neurons in awake mice while delivering a mild air puff (100 ms duration, ∼5 psi) to the ipsilateral eye (to avoid incidental stimulation of whiskers associated with the imaged neurons, Fig. 4B). In 15 of 21 L1 interneurons, the air puff evoked a clear depolarization. In 6 of these neurons the air puff evoked one or more spikes and in 3 of these neurons, the air puff evoked a barrage of firing that lasted ∼1 s, strikingly different from the precisely timed single spikes evoked by whisker stimulation (Fig. 4C). When the air puff was applied in the middle of an epoch of single cell-targeted optogenetic stimulation (500 ms, 5.8 mW/mm^2^), we did not observe a significant change in mean spike rate of the optogenetically targeted neurons (Fig. 4D). Some neurons increased their spike rate while others decreased their spike rate (see e.g. Fig. 4C, second and fourth traces).

**Figure 4.**
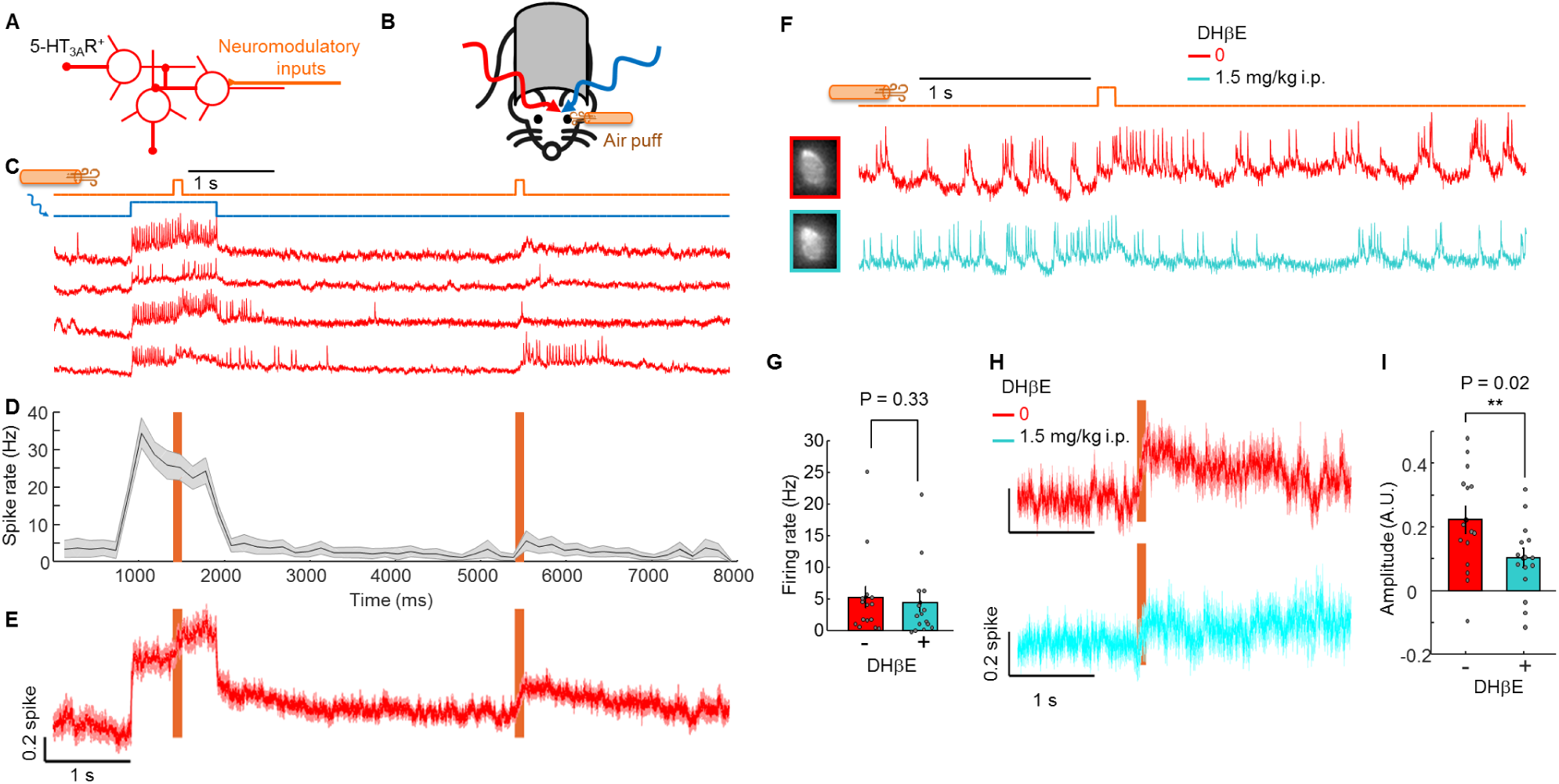
Cholinergic inputs drive excitation in L1 interneurons. (**A**) Schematic showing neuromodulatory inputs driving L1 interneurons. (**B**) Experiment to probe neuromodulatory effects in L1. Optopatch measurements were performed in barrel cortex L1 interneurons of awake 5-HT_3A_R-Cre mice while a mild air puff was applied to the ipsilateral eye. (**C**) Fluorescence recordings from single cells showing responses to air puff stimulation in the presence and absence of baseline optogenetic stimulation. (**D**) Mean spike rate during air puff stimulation with and without baseline optogenetic depolarization (*n* = 21 neurons, 4 mice). (**E**) Mean subthreshold response to air puff. Spikes were digitally removed from the traces. (**F**) Effect of a cholinergic blocker, DHβE, on the air puff response. Paired measurements were performed returning to the same cell before and after drug administration. (**G**) DHβE did not significantly affect spontaneous spike rate. (**H**) Mean subthreshold responses to air puff before and after administration of DHβE. (**I**) DHβE significantly reduced the subthreshold response to air puff, as quantified by the amplitude of the subthreshold response to air puff. Data in (G – I) from *n* = 16 neurons measured before and after drug administration, 3 mice.

To resolve the subthreshold dynamics, we digitally removed spikes and calculated the air puff-triggered average across all imaged neurons. In the absence of optogenetic stimulation, the air puff evoked a depolarization that grew over ∼100 ms, reached 20 ± 3% of spike height, and decayed with a ∼1600 ms recovery time (Fig. 4E). In the presence of single-cell targeted optogenetic stimulation, the air puff evoked a depolarizing transient in the subthreshold voltage of the optogenetically targeted cell (Fig. 4E), opposite to the hyperpolarizing transient evoked by a whisker stimulus under comparable conditions (e.g. Fig. 2C). Together, these results implied that the air puff evoked a predominantly excitatory input to the L1 microcircuit.

The α4 nicotinic acetylcholine receptor is highly expressed in L1 interneurons (*2, 31*), so we hypothesized that this receptor was mediating the neuromodulatory response. We made paired recordings of the same L1 interneurons before and after systemic administration of the α4 nAChR blocker dihydro-β-erythroidine hydrobromide (DHβE, 1.5 mg/kg i.p., Fig. 4F). This drug did not significantly affect the spontaneous spike rate (Fig. 4G), but it largely suppressed the air puff-induced depolarization, consistent with a cholinergic mechanism for this effect (amplitude, A. U., 0.22 ± 0.04 before vs 0.10 ± 0.03 after drug, *n* = 15 neurons, 3 mice, *p* = 0.02, two-sided paired-sample *t*-test, Fig. 4H, I).

### Numerical model of L1 microcircuit

Prior studies have characterized the anatomical and electrophysiological properties of L1 interneurons in detail (*2, 5, 14, 32*), and modeling efforts have led to anatomically and biophysically detailed simulations (*27, 33*). We sought to develop a numerical model of the L1 microcircuit that incorporated our data and prior information. Such a model should be able to reproduce our experimental results and to generate testable predictions for future experiments. Rather than aiming for numerical precision, we sought to build a simple model which captured the main features of the data in an intuitive and computationally efficient format.

A variety of classification schemes have been proposed for L1 neurons based on electrophysiology, morphology, or molecular markers (*11, 15*). Here we consider two broad classes based on firing properties and morphology. Elongated neurogliaform cells (eNGC) are slow to spike near threshold, do not adapt, and primarily synapse within L1 (*14, 15*). Single bouquet-like (SBC-like) cells burst readily, quickly adapt, and primarily dis-inhibit underlying cortex (*14, 15*). Both cell types receive thalamic and neuromodulatory inputs (*5*). Since the SBC-like cells do not synapse within L1, the dynamics can be split into the mutually inhibitory eNGC network, driven by thalamic and neuromodulatory inputs, and the SBC-like output, driven by the eNGC network, thalamic, and neuromodulatory inputs (Fig. 5). For present purposes we neglect cortico-cortical connections.

**Figure 5.**
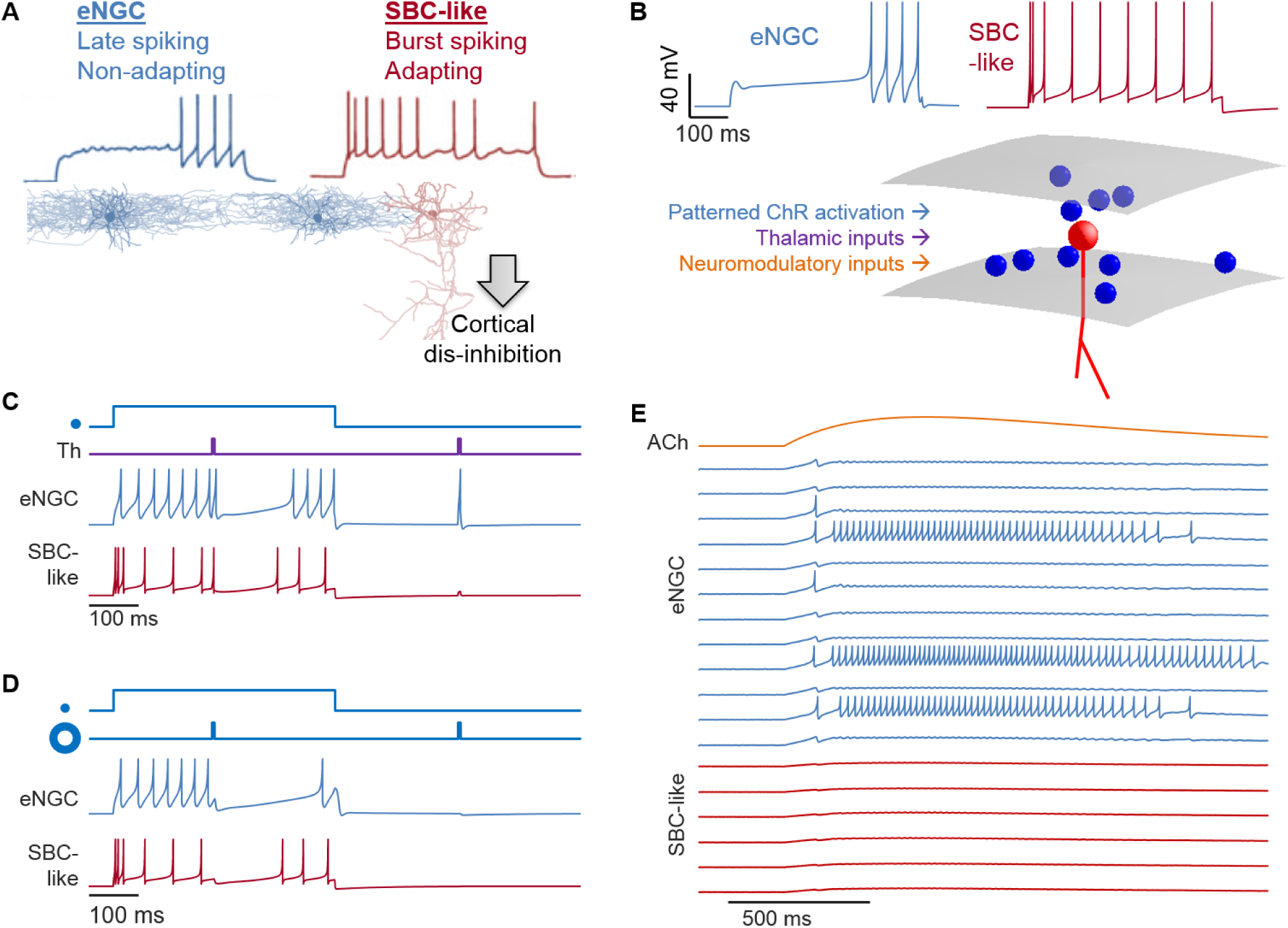
Computational model for the L1 micro-circuit. (A) Characteristic morphology and spiking patterns of L1 interneurons. Manual patch clamp recordings were acquired near rheobase. The eNGC cells form a mutually inhibitory network within L1. Downward-projecting SBC-like neurons receive inhibition from the eNGC network, and in turn dis-inhibit underlying cortex. Electrophysiology traces adapted from (Chu et al.J. *Neurosci* 23 (2003): 96-102.) and neuronal morphologies from (Jiang et al. *Science* 350 (2015): aac9462). (B) Model of L1 microcircuit. Neurons of both sub-types were randomly distributed within a disk-shaped barrel. Parameters of Izhikevich-type models were adjusted to mimic the characteristic firing patterns. Cells of both classes received optogenetic, thalamic, and neuromodulatory inputs. (C) Simulation of a complete L1 network for a single barrel (51 neurons). The network was exposed to network-wide thalamic excitation and single-cell targeted optogenetic stimulation. The traces show two simulations in which the optogenetic stimulation and voltage measurement were targeted either to an eNGC cell or to an SBC-like cell. Thalamic excitation alone induced spikes or sub-threshold excitation. In the presence of targeted optogenetic stimulation, thalamic excitation suppressed spiking. Compare to Figs. 2C, S8. (D) Simulation of an L1 network response to tonic optogenetic stimulation of a central neuron and pulsed optogenetic stimulation of the surrounding neurons. The traces show two simulations in which the central stimulation was targeted either to an eNGC cell or to an SBC-like cell. Surround stimulation hyperpolarized the targeted neurons and suppressed spiking. Compare to Fig. 3D. (E) Simulation of an L1 network response to a graded neuromodulatory input activating an excitatory conductance in all neurons. A few eNGC neurons spiked first, suppressing spiking of the rest of the network. The increased network inhibition blocked spiking of the output SBC-like neurons. The plots show a randomly selected subset of the 51 neurons in the circuit. Compare to Fig. 4C.

To capture the distinct firing properties of L1 interneurons, we used Izhikevich-type models (*34, 35*), with parameters tuned to match the observed spiking properties of eNGC and SBC-like interneurons (Fig. 5B, Methods, Table S1). Though not explicitly conductance-based, Izhikevich models can accommodate synaptic inputs as ohmic conductances (*36*), a critical feature for recapitulating the non-additivity of sensory and optogenetic stimuli. We used literature values to set the approximate density, length-scale, strength, and duration of synaptic connections (Methods, Table S2, Figs. S10-11). Detailed structural studies in rats have shown that each barrel contains ∼40-50 L1 interneurons, consistent with our estimates based on published micrographs from mice (*19*). The lateral extent of the axonal and dendritic trees is ∼200 – 250 µm, comparable to the size of the barrel. Consistent with this broad arborization, patch clamp measurements in slices found that most nearby pairs of L1 eNGC interneurons had synaptic connections (*15, 37*). We assumed that neurons did not synapse onto themselves. The time-dependent strengths of optogenetic, thalamic, and neuromodulatory inputs were varied to mimic different experimental conditions.

To characterize the single-cell models, we simulated individual eNGC and SBC-like neurons exposed to combinations of tonic excitatory and inhibitory inputs. Both cell types showed Type II firing patterns (*38*), i.e. abrupt onset of spiking during smoothly increased excitatory drive (Fig. S12). We simulated single-cell behavior under step-wise increasing optogenetic stimulation and with pre-specified transient excitatory (mimicking thalamic) and inhibitory (mimicking local network) inputs. The simulated cells produced spike patterns and subthreshold dynamics (Fig. S12) that resembled the corresponding experimental results (Figs. 2C, S8).

We then simulated a full L1 network for a single barrel: 51 L1 interneurons (34 eNGC, 17 SBC), randomly distributed in a region 300 µm on edge and 150 µm thick, with biologically plausible synaptic weights. Thalamic excitation and targeted optogenetic stimulation remained as input parameters, while inhibition was solely due to internal network dynamics. The simulated dynamics (Fig. 5C) closely matched the experimental results (Fig. 2C, S8). At baseline the model yielded sensory-evoked sub-threshold depolarization or single spikes. In the presence of optogenetic drive, the model yielded sensory-evoked single spikes followed by hyperpolarization and spike suppression. We then reproduced the donut stimulation experiments from Fig. 3 (Methods, Fig. S13). As anticipated, when a central neuron was optogenetically depolarized, optogenetic activation of surrounding neurons induced hyperpolarization and suppressed spiking of the central neuron (Fig. 5D).

The experimental responses to the air puff stimuli were highly heterogeneous (Fig. 4C), an effect which we initially ascribed to variability in neuron sub-types or to specific patterns of connectivity not included in our model. Despite the overly simplistic model, we studied how the simulated network responded to neuromodulatory inputs. We modeled neuromodulation as a gradual activation of an excitatory conductance in all L1 interneurons (Methods). To our surprise, the simulations of a nearly homogeneous eNGC network recapitulated the highly heterogeneous responses observed experimentally (Fig. 5E), and even captured the approximate proportions of cells showing subthreshold depolarizations, isolated spikes, and tonic firing.

This surprising symmetry breaking (emergence of qualitatively distinct responses from a nearly homogeneous population) was explained by the mutually inhibitory nature of the L1 eNGC network. A few neurons, by chance, spiked first. These drove inhibition in their neighbors, suppressing spiking in response to the tonic excitation. Thus the highly heterogeneous single-cell responses to neuromodulatory inputs can be explained by an emergent network phenomenon and do not require sub-populations with specific wiring or electrophysiological properties (though our experiments do not inform whether functionally distinct sub-populations contribute to the heterogeneous responses *in vivo*).

Based on the success of the simple model, we suggest an intuitive picture for how sensory and neuromodulatory inputs interact in L1. We propose that a transient thalamic input initially activates the output (SBC-like) cells, while a sustained thalamic input predominantly drives eNGC-mediated inhibition and suppresses the output. Numerical simulations showed that this is the case (Fig. S14). Within this picture, the distinct intrinsic firing properties of the different L1 sub-types play a crucial role in tuning the circuit as a novelty detector. Our model predicts that under weak neuromodulatory excitation, the network becomes sensitized to thalamic inputs because all cells are closer to threshold; but under strong neuromodulatory excitation the emergence of sustained spiking in a subset of eNGC cells may suppress network responses to thalamic inputs. This is a testable prediction.

The L1 microcircuit is particularly suited for computational modeling due to the relative simplicity of the internal connectivity, the absence of recurrent excitation, and the sparse distributions of cells. Our computationally efficient model is readily applied to large-scale simulations. These simulations could be used to make testable predictions of how the L1 microcircuit would respond to various combinations of sensory and neuromodulatory inputs, patterned in space and time. While our simple model lacks many features of the real circuit, we suggest that this model is a useful starting point from which to add more realism.

## Discussion

Our experiments revealed that lateral inhibition among L1 interneurons mediates precisely timed single-spike responses to abrupt sensory inputs. A striking aspect of these findings was that the same sensory input (a whisker deflection) could depolarize and elicit spikes in a hyperpolarized neuron, but hyperpolarize and suppress spikes in a depolarized neuron. This observation highlights the importance of considering electrophysiological context when interpreting functional data recorded *in vivo*.

Remarkably, this highly nonlinear effect (opposite sign response depending upon initial membrane voltage), was quantitatively explained by a simple biophysical model which only contained idealized batteries and linear, ohmic conductances. The appearance of nonlinear responses in a linear membrane model is explained by the fact that, even in a linear model, the voltage is a nonlinear function of the conductances (see Supplementary Text). The oft-used approximation that excitatory and inhibitory inputs simply add to make a net synaptic current is clearly violated.

While our experiments used a channelrhodopsin to drive depolarization, ionotropic AMPA receptors (*39*), acetylcholine receptors (*40*), serotonin receptors (*41*) and channelrhodopsins (*42*) all have similar current-voltage relations, implying that baseline activation of any of these receptors could switch the sensory-evoked response of an L1 neuron from excitation-dominated to inhibition-dominated.

Our results further showed that thalamic and neuromodulatory excitation converge in L1 neurons, albeit with different temporal profiles and consequently different network effects. Whereas rapid thalamic excitation led to synchronous spiking followed by network inhibition, slower neuromodulatory excitation drove tonic spiking in a subset of cells which suppressed spiking in the majority.

Experiments *in vivo* (*3*) and in slices (*43*) have shown that cholinergic stimulation elicits complex effects on L1 interneurons, but how these effects modulate sensory-evoked responses *in vivo* remains to be determined. A clear goal for future work will be to characterize the role that each subclass of L1 interneurons plays in integration of sensory, neuromodulatory, and cortico-cortical inputs.

All-optical electrophysiology can report both the nature of the synaptic inputs (E vs. I) and the spiking output of a cell, revealing the transformation that the cell implements. Optogenetic stimulation and voltage imaging in distinct neural populations can reveal cell type-specific connections and their role in circuit dynamics. While we focused on rapid sensory processing, these tools may also prove useful in studies of neural plasticity, development and disease mechanisms.

## Acknowledgments

We thank B. Sabatini for advice and discussion; G. Vargish and T. Hensch for the 5HT_3A_R-Cre mouse line; S. Begum, K. Williams, A. Preecha and H. Dahche for technical assistance; M. Gomez-Ramirez, C. Moore and F. Wang for advice on whisker stimulation; H. Pi for advice on air puff; B. Gmeiner for advice on optics; Y. Adam for advice on mouse surgeries.

## Author contributions

L.Z.F., A.E.T. and A.E.C. conceived and designed the study. L.Z.F. designed and conducted the experiments, built the optical system, programed the software, analyzed the data and co-wrote the manuscript. S.K. and H.W. built the optical system in early stage. U.B. performed numerical simulations and data fitting. K.D.P. and E.S.B. shared SomArchon. V.P. helped with slice experiments. M.E.X. helped analyze the data. A.E.C. performed network simulations, co-wrote the manuscript and supervised the research.

## Data and materials availability

Plasmids are available on Addgene. Data, code, and materials used in the analysis are available upon reasonable request to A.E.C..

## Supplementary Materials

Materials and Methods

Supplementary Text

Figs. S1 to S14

Tables S1-S3

