## Supplementary materials for "All-optical electrophysiology reveals excitation, inhibition, and neuromodulation in cortical layer 1"

**This PDF file includes:**

Materials and Methods
Supplementary Text
Figs. S1 to S14
Tables S1-S3

### Materials and Methods

#### Design of Optopatch4

Optopatch4 construct (SomArchon-eGFP-P2A-somCheRiff) was cloned into an AAV vector with Cre-dependent expression driven by the hSyn promoter.

LZF1735 pAAV_hSyn-DiO-SomArchon-eGFP-P2A-somCheRiff (Addgene #126512)

LZF1733 pAAV_CAG-FLEX-SomArchon-eGFP

High-titer AAV2/9 virus with Optopatch4 ($1.74\times10^{13}$ GC/mL) was obtained from the Janelia Farm Vector Core. High-titer AAV2 virus with LZF1733 ($6.30\times10^{12}$ GC/mL) was obtained from the UNC Vector Core. High-titer AAV9 virus with CKII(0.4)-Cre ($2.8\times10^{13}$ GC/mL) was obtained from UPenn Vector Core.

#### Optical system for holographically targeted voltage imaging and patterned optogenetic stimulation

The optical system combined a red laser ($\lambda$ = 639 nm) path for holographic targeted illumination voltage imaging, a blue laser ($\lambda$ = 488 nm) path for micromirror-patterned optogenetic stimulation, a two-photon (2P) path for structural imaging, and a wide-field epifluorescence imaging path.

*Red laser path.* A red laser (CNI Inc., MRL-FN-639, $\lambda$ = 639 nm, 700 mW single transverse mode) was coupled into the setup via a photonic crystal polarization maintaining fiber (NKT Photonics, LMA-PM-15). The fiber output was collimated with an $f$ = 100 mm focal length lens (Thorlabs, AC254-100-A-ML) to form a beam with approximately ~10 mm diameter. The polarization of the beam was set with a zero-order half-wave plate. The beam was directed onto a holographic reflection-mode liquid crystal spatial light modulator (SLM, Meadowlark 1920SLM VIS) with a resolution of 1920 x 1152 pixels. Zero-order diffraction was blocked by a home-made anti-pinhole comprised of a dot of solder on a glass slide, placed in a plane conjugate to the sample image plane. The SLM was re-imaged onto the back-focal plane of the objective via a series of relay optics. The objective lens was a 25× water immersion objective, numerical aperture 1.05 (Olympus XLPLN25XWMP2). A mechanical shutter blocked the red laser between data acquisitions. A series of OD filters were placed after the red laser for modulating intensity.

In the first generation of the setup, we used a variable focal length camera lens (Sigma macro 18-200 mm) to control the magnification of the SLM at the back focal plane of the objective. Demagnifying the SLM decreased the effective numerical aperture of the illumination at the sample, leading to bigger spots in the sample, but also to a larger region that could be targeted with red light. In the second-generation system, we used a fixed lens after the SLM to minimize aberrations. All relay lenses are specified in Table S3.

The SLM device was controlled by custom software. A user specified a set of lines for the SLM to target by drawing on a wide-field epifluorescence image or a 2P fluorescence image.

These lines were discretized into a set of spots. The SLM phased pattern was calculated using the Gerchberg-Saxton algorithm.

Red laser intensity was ~ 3 mW per cell for *in vivo* imaging, ~ 1 mW per cell for acute slice imaging.

Blue laser path. A blue laser (Cobolt, 06-01 series, $\lambda$ = 488 nm, 60 mW) was modulated in intensity via an acousto-optic tunable filter (AOTF; Gooch and Housego TF525-250-6-3-GH18A). The beam was focused into a single-mode optical fiber. The output was collimated with an $f$ = 60 mm focal length lens (Thorlabs, AC254-060-A-ML) to form a beam with approximately a ~17 mm diameter. The beam was then sent to a digital micromirror device with a resolution of 1024 x 768 pixels (DMD, Vialux, V-7001 VIS). The patterned blue beam was combined with the patterned red beam via a dichroic mirror. The DMD was re-imaged onto the sample at a magnification such that one DMD pixel corresponded to 0.62 μm in the sample plane. The DMD optical system enabled patterned blue light stimulation across a field of view of ~450 x ~520 μm.

The DMD was controlled by custom software. For excitability measurement, a pixel bitmap was preloaded onto and projected from the DMD. For lateral inhibition experiments, pixel bitmaps were loaded into the on-board RAM and digital clock pulses triggered the DMD to sequence through the pre-defined set of exposure patterns.

Wide-field fluorescence imaging path. The image was relayed from the objective to the camera via a series of three lenses. The final image formation step was performed by a 4x objective (Olympus XLFLUOR 4X/340) serving the role of the tube lens. Fluorescence was collected on a scientific CMOS camera (Hamamatsu ORCA-Flash 4.0). The final magnification of the optical system was 16.7, corresponding to 0.39 μm in the sample plane per camera pixel.

Fluorescence from the sample was separated from the blue and red excitation beams via a dichroic mirror (Di03-R405/488/561/635-t3-40x55). An emission filter (Semrock 635 nm long-pass, BLP01-635R-25) further separated SomArchon fluorescence from scattered excitation light. An IR-blocking emission filter (Semrock, FF01-842/SP-25) was placed for blocking scattered infrared excitation light.

All movies are acquired at 1 kHz. To image at 1 kHz, the camera region of interest (ROI) was restricted to typically 200 rows, centered on the image-sensor midline.

The imaging system was designed for a magnification lower than the nominal 25x of the objective for two reasons. First, lower magnification increased the number of neurons that could be imaged simultaneously onto the limited detector area accessible at 1 kHz. Second, by concentrating sample photons onto as few camera pixels as possible, we sought to minimize the contribution from camera electronic noise, so that all signals would be in the shot noise-limited regime.

*Two-photon imaging path*. Light from a femtosecond tunable pulsed infrared laser (Spectra Physics DeepSee) was sent to a pair of galvo mirrors (Cambridge Technologies 6215H). The galvos were re-imaged onto the back focal plane of the objective via an optimized scan lens (Thorlabs, SL50-CLS2) and tube lens (Thorlabs, TL200-CLS2). The visible (blue and red) and near-infrared beams were combined using a 785 nm long-pass dichroic mirror (Semrock, Di03-R785-t3-40x55). GFP fluorescence was directed to the 2P detection path via a removable 550 nm short-pass dichroic. Scattered excitation light was blocked by a 633 nm short-pass emission filter, an IR-blocking emission filter (BSP01-785R-25) and a band-pass emission filter (FF03-525/50-25). A pair of lenses (focal lengths 75 mm and 16 mm) re-imaged the back-aperture of the objective onto a photomultiplier tube (Hamamatsu, H11706P-40). The output of the photomultiplier was amplified and low pass filtered through an amplifier unit (Hamamatsu C7319) and then digitized.

*Control software*. The entire setup was controlled by custom software written in LabView. Interfacing was via a National Instruments DAQ (NI PCIe-6363).

The software contained routines for registration of the DMD, SLM, 2P microscope coordinates to the camera via affine transformations. The camera served as the global reference coordinate system.

Experimental protocols were specified by a set of images (to the SLM and the DMD), output waveforms (to the galvos, the AOTF, the shutters, the update clock on the DMD, the piezo whisker stimulator, and the air puff controller), and analog input streams (from the PMT, the camera exposure clock, and a patch clamp electrophysiology setup not used in the present work).

The Hamamatsu camera uses an internal 100 kHz clock to synchronize image row readout. We found that when the camera exposure times were triggered by the DAQ in synchronous mode, the camera rounded the exposure time to the nearest 10 μs, leading to 1% jitter in exposure time for 1 kHz imaging. To address this noise source, we used a custom firmware upgrade to access the 100 kHz camera clock. This clock became the master clock for the DAQ system, and all analog and digital input/output functions were synchronized to the camera clock.

**Imaging in acute slices**

All procedures involving animals were in accordance with the National Institutes of Health Guide for the care and use of laboratory animals and were approved by the Institutional Animal Care and Use Committee at Harvard University.

*Virus injection for acute slice measurements.* Virus comprising AAV2/9 hSyn-Dio-SomArchon-eGFP-P2A-somCheRiff ($1.74\times10^{13}$ GC/mL) was diluted in PBS and injected at a final titer of ~$2\times10^{12}$ GC/mL.

5HT3AR-Cre$^{+/-}$ mice were crossed with wild-type C57BL/6 mice. Pups were cryo-anesthetized at P0-P2 and immobilized dorsal side up under a stereotaxic microscope. Injections were made using home-pulled micropipettes (Sutter P1000 pipette puller), mounted in a microinjection pump (World Precision Instruments Nanoliter 2010) controlled by a microsyringe pump controller (World Precision Instruments Micro4). The micropipette was positioned using a stereotaxic instrument (Stoelting Digital Mouse Stereotaxic Instrument). Pups were injected in the left hemisphere, 1 mm lateral and 1.2 mm anterior to lambda. Starting at a depth of 0.3 mm beneath the surface of the skull, virus injections (40 nL, 1 nL/s) were performed at 0.1 mm increments as the pipette was withdrawn. Pups were placed back in their home cage once they were awake.

*Genotyping.* Genotyping for $5HT_{3A}R$ was performed with the PCR primer pairs: Cre 5': 5' TAT CTC ACG TAC TGA CGG TG 3' and Cre 3': 5' AGA CTA ATC GCC ATC TTC CAG C 3' to yield a 500 bp band from Cre.

*Acute slice preparation.* Acute brain slices were prepared from P16–P28 $5HT_{3A}R\text{-}Cre^{+/-}$ mice. The mice were anesthetized by isoflurane and then perfused with carbogen (95% $O_2$, 5% $CO_2$)-saturated ice-cold slicing solution with the following composition (in mM): 110 choline chloride, 2.5 KCl, 1.25 $NaH_2PO_4$, 25 $NaHCO_3$, 25 glucose, 0.5 $CaCl_2$, 7 $MgCl_2$, 11.6 Na-ascorbate, and 3.1 Na-pyruvate. Mice were then decapitated and the brains were rapidly coronally sliced with 300 µm thickness on a vibratome (Leica VT 1200S).

Slices were incubated for 45 min at 34 °C in a carbogenated artificial CSF (ACSF) with the following composition (in mM): 127 NaCl, 2.5 KCl, 1.25 $NaH_2PO_4$, 25 $NaHCO_3$, 25 glucose, 2 $CaCl_2$, and 1 $MgCl_2$. The osmolarity of all solutions was adjusted to 300–310 mOsm and the pH was maintained at 7.3 under constant bubbling with carbogen.

*Imaging acute slices.* Measurements were conducted in ACSF at 23 °C under ambient atmosphere. The slice was immobilized in a slice recording chamber using a slice anchor (Warner Instruments, SHD-40/2). ACSF, perfused with carbogen, was flowed through the chamber at a rate of 2 mL/minute.

**Cranial windows and virus injections**

*Cranial window surgery and virus injection for imaging barrel cortex L1.* Virus comprising AAV2/9 hSyn-Dio-SomArchon-eGFP-P2A-somCheRiff ($1.74\times10^{13}$ GC/mL) was diluted in PBS and injected at a final titer of ~$3\times10^{12}$ GC/mL.

The procedure for surgery and imaging in barrel cortex L1 followed the protocol from Andermann (*1*). 35-60-day-old heterozygous $5HT_{3A}R$-Cre mice (male and female) were deeply anesthetized with 2% isoflurane and maintained with ~1% isoflurane throughout the surgery. Eyes were kept moist using ophthalmic eye ointment. Body temperature was continuously monitored and maintained at 37 °C using a heating pad (WPI, ATC2000). The skull was exposed and thoroughly dried and a 3 mm round craniotomy (3.3 – 3.4 mm lateral, 1.6 mm caudal of bregma) was opened using a biopsy punch (Miltex). Virus was then injected in 4 - 8 locations in the center of the craniotomy. Starting at a depth of 0.2 mm beneath the surface of the dura, virus injections (60 nL, 1 nL/s) were performed at 0.1 mm increments as the pipette was withdrawn. Brain surface was kept moist with saline throughout the injection.

A window was prepared prior to the surgery and comprised two 3 mm round #1 cover glasses and one 5 mm round #1 cover glass (Harvard apparatus) cured together with UV curable adhesive (Norland Products, NOA 81). Following the virus injection, the window was then placed covering the barrel cortex and cemented to the skull with dental cement (C&B metabond, Parkell, No. 242-3200). After the window cured, a titanium headplate (similar to the design in Ref. (*1*)) was glued around the window and any exposed skull was covered with dental cement. Animals were returned to their home cage for recovery and treated for 3 days with Carprofen (5 mg/kg) and Buprenorphine (0.1 mg/kg) twice a day. To avoid damage to the implant, mice were housed in separate cages.

*Cranial window surgery and virus injection for L1-3 imaging* (Fig. S4). Virus comprised AAV2 CAG-FLEX-SomArchon-eGFP (final titer ~$0.5\times10^{12}$ GC/mL) mixed with CKII(0.4)-Cre virus (UPenn vector core, final titer ~$1\times10^{11}$ GC/mL).

The procedure for surgery and imaging in visual cortical L1-3 followed the same protocol as above, except that 35-60-day-old wild-type C57BL/6 mice (male and female) were used. The coordinates for the 3 mm round craniotomy were 2.4 mm lateral and 2.7 mm caudal of bregma. Virus was then injected in 4 - 8 locations in the center of the craniotomy. Starting at a depth of 0.3 mm beneath the surface of the dura, virus injections (60 nL, 1 nL/s) were performed at 0.2 mm increments as the pipette was withdrawn.

**Tracking, whisker stimulation, intrinsic imaging and *in vivo* voltage imaging**

An IR LED light (850 nm) was placed in front of the animal. A PointGrey camera (GS3-U3-51S5M-C, Mono Grasshopper3 USB 3.0 Camera) with a Fuji lens (Fuji Photo Optical 1:1.4/25 Fujinon-TV Camera Lens) and an IR-passing optical filter (Thorlabs, FB850-40) was placed on the side to track the animal's face (whisker motion and eye blinks) during data acquisitions.

*Whisker stimulation.* An individual whisker (typically B2, C2 or D2; other whiskers were trimmed to prevent direct contact with the glass pipette) was inserted into a glass pipette glued to a piezoelectric actuator. The actuator was connected to an amplifier (Krohn-Hite 7602M) and controlled by the DAQ.

*Intrinsic imaging.* Intrinsic imaging was performed on the same setup as described above. A 4x objective (Olympus XLFLUOR 4X/340) was used to image the entire 3 mm cranial window. The whisker stimulation was 10 Hz for 4 s with a 16 s interstimulus interval. A red LED (625 nm) illuminated the window surface from the side. Reflected light was imaged onto the camera at 10 Hz frame rate. A decrease in reflectance from the brain indicated the barrel, which could be localized relative to the blood vessel pattern as visualized with 488 nm illumination.

*Imaging anesthetized animals*. Imaging started 3 weeks post-surgery. Mice were lightly anesthetized (0.7–1% isoflurane), head-fixed under the upright microscope using the titanium head plate and held in a body tube. Eyes were kept moist using ophthalmic eye ointment. Body temperature was continuously monitored and maintained at 37 $^{o}$C using a heating pad (WPI, ATC2000). A typical imaging session lasted 1–2 hours, and then animals quickly recovered and returned to their home cage. Recordings targeting L1 neurons *in vivo* were performed at a depth < 150 μm in both anesthetized and awake animals.

*Habituation and imaging awake animals.* Habituation started 2 weeks post-surgery. Each animal was acclimated to the head restraint in a body tube for at least 3 days before starting the imaging sessions. For imaging in awake animals, a 3D-printed paw blocker was placed in front of the forepaws to prevent them from pushing away the glass pipette for whisker stimulation.

*Air puff.* The timing of the air puff was controlled by a solenoid valve (WPI). The strength of the air puff was set ~ 5 psi and controlled by a pressure regulator (Festo, pressure regulator LRP-1/4-4). The air puff was delivered through a blunt needle at ~ 5 mm from the eye ipsilateral to the brain hemisphere used for voltage imaging (to avoid spurious whisker stimulation arriving in the imaged barrels). Air pressure and needle position were adjusted to achieve an air puff strength just strong enough to evoke an eye blink response and increase the pupil diameter.

*DHβE administration.* To modulate cholinergic signaling, dihydro-β-erythroidine hydrobromide (DHβE, Tocris; 2349) was diluted in saline. The drug was administered systemically (1.5 mg/kg i.p.). Optopatch and air puff measurements were performed before, and then 30 min. after drug administration, on the same sets of cells. The experiment was performed on each animal twice, on successive days.

**Center/surround optogenetic stimulation**

For lateral inhibition experiments, we defined two optogenetic stimulus patterns. "Central masks" covered individually the cell bodies of 1 – 3 neurons at the center of the field of view. We calculated the geometrical centers of these masks individually. The mean of these centers was set as the coordinates for the center for the surrounding "annulus mask".

The annulus inner radius was set to be ~100 μm from the most non-centered central mask. This distance was selected to minimize the impact of scattered light from the annulus mask. The outer radius of the annulus mask was set as the largest value at which the annulus would be contained within the FOV. Typical outer radii were ~200 μm.

The image sequence was composed of three composite masks: (1) central masks only; (2) central mask and annulus mask together; (3) annulus mask only. These masks were preloaded into the on-board RAM and digital clock pulses triggered the DMD to sequence through the pre-defined set of exposure patterns.

**Data analysis**

Data were analyzed with homemade code written in MATLAB.

*Corrections for photobleaching and motion artifacts.* Movies were first corrected for motion using the NoRMCorre algorithm (*2*). Movies were then corrected for photobleaching by dividing the movie by an exponential fit of the mean fluorescence.

*Image segmentation and waveform extraction.* We divided the movie into sub-movies based on patterns of illumination from the DMD masks and performed activity-based image segmentation separately in each sub-movie. Whereas subthreshold voltages could be correlated between a cell and out-of-focus background cells, we assumed that spiking was not correlated with background, and furthermore that the spatial footprint associated with spiking would be the same as for true subthreshold dynamics. To remove subthreshold signals for segmentation purposes, movies were filtered in time with a 100 Hz high-pass filter. Movies were then segmented semi-automatically using principal components analysis followed by time-domain independent components analysis (PCA/ICA) (*3*). The spatial masks from PCA/ICA were then applied to the original movies without high-pass filtering to extract fluorescence traces.

*Removing scattering background for lateral inhibition measurements.* Background fluorescence from the region surrounding the central imaged cells (due to the scattered light) was subtracted from the baseline fluorescence of the cell.

*Spike finding and scaling of fluorescence recordings.* A simple threshold-and-maximum procedure was applied for spike detection. Fluorescence traces were first high-pass filtered, and initial threshold was set at 3 times the noise level. This threshold was then manually adjusted if needed.

All fluorescence signals were normalized to spike height for spike triggered average or stimulation triggered average.

*Spike removal for calculation of subthreshold waveforms.* Spikes were digitally removed and replaced with interpolations of the surrounding data. Spike width was estimated by viewing individual fluorescence recordings. Linear interpolations were performed between data-points 1 ms beyond the edges of the spike.

**Statistics**

All error ranges represent standard error of the mean, unless otherwise specified. For the same neurons before and after drug administration, and in anesthetized and awake states, paired sample t-test was used. For two-sample comparisons of a single variable, student's t-test was used. Probabilities of the null hypothesis $p < 0.05$ were judged to be statistically significant.

**Biophysical modeling of membrane potential**

The evolution of membrane potential in the presence of synaptic inputs and optogenetic stimulation (Fig. 2) was simulated with a passive single compartment model using the following equation:

$$C_m \frac{dV}{dt} = g_e(E_e - V) + g_i(E_i - V) + g_l(E_l - V) + g_{ChR}(E_{ChR} - V), \qquad (1)$$

where $C_m$ is the membrane capacitance, $g_e$, $g_i$, $g_l$ and $g_{ChR}$ are the conductance of excitatory, inhibitory, leak and channelrhodopsin channels, and $E_e$, $E_i$, $E_l$ and $E_{ChR}$ are the respective reversal potentials. The time course of conductance upon excitatory or inhibitory synaptic input was simulated using an alpha function:

$$g(t) = \begin{cases} g_{\text{syn}} \frac{t-t_0}{\tau} e^{1-\frac{t-t_0}{\tau}} + g_{\text{baseline}} & \text{for } t \geq t_0 \\ g_{\text{baseline}} & \text{for } t < t_0 \end{cases}. \qquad (2)$$

Here $g_{syn}$ is the strength of the synaptic input, $t_0$ is the time of the synaptic input, t is a time-constant of synaptic input, and $g_{baseline}$ reflects the tonic level of synaptic input excluding the event of interest.

In principle, the values of $g_{baseline}$ for the inhibitory and excitatory synaptic inputs could be wrapped into the definitions of $g_l$ and $E_l$. Doing so would not affect the solutions to Eq. 1. We chose to keep the baseline synaptic conductances as separate parameters to facilitate explorations of the model under different brain states (e.g. anesthesia vs. wakefulness). In this approach, $g_l$ and $E_l$ reflect cell-autonomous leak conductances (e.g. $K_{ir}$ channels), assumed to be independent of brain state, while $g_{baseline}$ captures the effect of network-dependent inputs.

To simulate lateral inhibition in L1, we assumed inhibition lagged excitation by 2 ms. Other parameters are listed below:

| Parameter | Value |
|---|---|
| $g_{e\,syn}$ | 1.5 nS |
| $g_{e\,baseline}$ | 0.1 nS |
| $g_{i\,syn}$ | 5 nS |
| $g_{i\,baseline}$ | 0.1 nS |
| $g_l$ | 3.33 nS |
| $g_{ChR}$ | 0 − 10 nS |
| $E_e$ | −5 mV |
| $E_i$ | −70 mV |
| $E_l$ | −70 mV |
| $E_{ChR}$ | 0 mV |
| $C_m$ | 150 pF |
| $\tau$ | 1 ms |

Equation 1 was numerically integrated using Euler's method.

The model above can be solved analytically for the steady-state voltage by setting $\frac{dV}{dt} = 0$, which yields:

$$V = \frac{E_e g_e + E_i g_i + E_l g_l + E_{ChR} g_{ChR}}{g_e + g_i + g_l + g_{ChR}}. \qquad (3)$$

The PSP amplitudes are obtained by calculating the difference in steady-state voltage, $\Delta V_{PSP}$, after vs. before the sensory perturbations to $g_e$ and $g_i$, assuming that all other parameters do not vary during the synaptic event. If one assumes that $g_e$ and $g_i$ are both zero before the synaptic event (i.e. by absorbing the pre-stimulus values of $g_e$ and $g_i$ into the definition of $g_l$ and $E_l$), then one finds:

$$\Delta V_{IPSP} \approx \frac{g_{ChR}[g_e(E_e - E_{ChR}) + g_i(E_i - E_{ChR})] + g_l g_e(E_e - E_l) + g_l g_i(E_i - E_l)}{(g_l + g_{ChR})(g_i + g_l + g_{ChR})}. \qquad (4)$$

Since only differences in voltage appear in Eq. 4, one can arbitrarily choose to set one of the voltages to zero, and measure all other voltages relative to this reference. For convenience, we set $E_l$ to zero for the fitting. Eq. 4 is linear in $g_{ChR}$ in the numerator, and quadratic in $g_{ChR}$ in the denominator, suggesting 5 fitting parameters to specify the function $\Delta V_{IPSP}(g_{ChR})$. However, the proportionality between $\Delta V$ and $\Delta F$ is not *a priori* known. Also the proportionality between $g_{ChR}$ and $I_{488}$ is not *a priori* known. Thus the shape of the function $\Delta F_{PSP}(I_{488})$ is governed by four fitting parameters.

To estimate the waveforms of the post-synaptic potentials, we first normalized all fluorescence traces by spike height. We then calculated the mean fluorescence over 10 ms before the whisker stimulus, and the mean fluorescence over 10 ms starting 30 ms after the whisker stimulus. The difference between these values was taken as the amplitude of the post-synaptic potential. Equation 4 was fitted to the data using the nonlinear least-squares method in Matlab.

*Analytical approximation.* To gain an intuition for the responses, one can make a simple estimate of the amplitude of the IPSP by assuming that $g_e = 0$ at all times, that $g_i = 0$ before the synaptic input, and that the resting potential is the same as the inhibitory reversal potential, i.e. $E_l \approx E_i$. One then obtains:

$$\Delta V_{IPSP} \approx \frac{g_i g_{ChR}(E_l - E_{ChR})}{(g_l + g_{ChR})(g_i + g_l + g_{ChR})}. \qquad (5)$$

Within the blue light intensity range used in our experiments, the CheRiff conductance is well approximated by a linear function of the blue intensity, though the proportionality factor depends on the (unknown) CheRiff expression level and attenuation of the blue light by scattering.

Eq. 5 shows that for small $g_{ChR}$ the amplitude $\Delta V_{IPSP}$ is linear in $g_{ChR}$, while for large $g_{ChR}$ the quadratic term in the denominator dominates and $\Delta V_{IPSP}$ decreases inversely with $g_{ChR}$. The value of $g_{ChR}$ that gives the largest amplitude IPSP is $g_{ChR}^{max} = \sqrt{g_l(g_l + g_i)}$, or for weak inhibition, $g_{ChR}^{max} \approx g_l$. If $g_l$ is large, then the shunting from $g_{ChR}$ is suppressed. The membrane time constant is approximately $\tau \approx \frac{C}{g_l + g_{ChR}}$. The inverse relation between $\tau$ and $g_{ChR}$ is consistent with our observation of faster recovery at stronger stimulus strength.

### Numerical model of L1 dynamics

*Single-cell properties.*

We simulated dynamics of L1 interneurons using Izhikevich-type models (*4*). We judged more detailed channel-based biophysical models to have too many unknown parameters and to be too computationally expensive for facile exploration of multiple conditions. Simpler spike rate-based models did not capture the details of the spike timing which we judged important for L1 circuit function.

We simulated the following dynamics for eNGC cells:

$$C\frac{dV}{dt} = k(V - V_r)(V - V_t) - u + I + G_{ds}(V_D - V) + g_{Exc}(V_{Exc} - V) + g_{Inh}(V_{Inh} - V) \quad (6)$$

$$\frac{dV_D}{dt} = G_{SD}(V - V_D)$$

$$\frac{du}{dt} = a(b(V - V_r) - u)$$

If $V > V_t$ then

$V \leftarrow c$

$u \leftarrow u + d$

The dynamics of SBC-like cells followed the same equations, but without the dendritic compartment ($G_{SD} = G_{DS} = 0$, $V_D = 0$). The meanings and values of the parameters for the two cell types are given in Table S1. The *a*, *b*, *c*, and *d* parameters were randomized by 10% between cells to prevent numerical degeneracies.

The differential equations were integrated using the Euler method with a step size of 0.1 ms. Some ancillary results are useful in tuning the properties of this model. These are:

1) In the absence of synaptic inputs ($g_{Exc} = g_{Inh} = 0$) or injected current ($I$ = 0), the resting-state membrane resistance is:

$$\left.\frac{dV}{dI}\right|_{I=0} = (b + k(V_t - V_r)).$$

2) To change the membrane resistance while maintaining dynamical properties, one should scale *b* and *k* proportionally.
3) To change the membrane capacitance while maintaining dynamical properties, one should scale *C*, *b*, *d*, and *k* proportionally.

*Channelrhodopsin activation.*

The channelrhodopsin CheRiff was modeled as an excitatory conductance with reversal potential 0 mV and conductance proportional to blue light illumination intensity. Channel gating kinetics were assumed to be instantaneous. Patterns of blue light were targeted to one or more cells and modulated in time during the simulation.

*Synaptic properties: Inhibition.*

Examination of patch clamp recordings of inhibitory post-synaptic potentials (IPSPs) in acute slices showed a rapid onset followed by a slow recovery. The recovery was not well captured by a single exponential. To approximate these dynamics we used the following function:

$$g_{inh}(t) = g_{inh}^{0}\left(\frac{t}{\tau_1}e^{1-\frac{t}{\tau_1}} + 0.6\frac{t}{\tau_2}e^{1-\frac{t}{\tau_2}}\right)$$

$\tau_1$ = 7 ms, $\tau_2$ = 35 ms. We found that the qualitative network dynamics were insensitive to variations in the functional form or time constants. One may think of the two terms as representing $GABA_A$ and $GABA_B$ receptors respectively, though a more accurate implementation of a $GABA_B$-mediated hyperpolarization would use the $K^+$ reversal potential (-90 mV) rather than the $Cl^-$ reversal potential (-70 mV).

To calibrate the value of $g_{inh}^{0}$ for eNGC → eNGC and eNGC → SBC synapses, we set up a model circuit with one eNGC cell synapsing onto an eNGC cell and an SBC cell. In the simulations, we adjusted a channelrhodopsin conductance ($V_{ChR}$ = 0 mV) to depolarize the downstream cells to -55 mV. We then triggered the upstream cell to spike. We adjusted $g_{inh}^{0}$ to induce IPSP amplitudes of -1.5 to -2 mV, to match the patch clamp data. The model did not contain short term inhibitory synaptic plasticity.

The IPSP amplitudes recorded via patch clamp were assumed to represent the strongest possible IPSPs. The IPSP strength between each pair of neurons in the circuit was modulated by a Gaussian function of separation, with a length-scale set by the sum of sizes of the presynaptic axonal arbor and the postsynaptic dendritic arbor.

*Synaptic properties: Excitation.*

We modeled thalmocortical excitation in L1 using the function:

$$g_{exc}(t) = g_{exc}^{0}\frac{t}{\tau}e^{1-\frac{t}{\tau}}$$

We calibrated the amplitude of $g_{exc}^{0}$ and $\tau$ by simulating thalamic inputs to eNGC cells and SBC-like cells and matching to literature data which showed that whisker-evoked EPSP amplitudes *in vivo* were 3 – 7 mV. We assumed that the timecourse of thalamic excitation to all neurons was identical. Excitatory synaptic strengths were randomized by 20% between cells to prevent numerical degeneracies. The model did not contain short term excitatory synaptic plasticity.

*Synaptic properties: Neuromodulation.*

The greatest uncertainty in the model surrounded the timecourse and strength of neuromodulatory inputs. We assumed that neuromodulatory inputs activated an excitatory conductance with reversal potential 0 mV. We assumed that the strength and timecourse of neuromodulatory action was the same on the eNGC and SBC-like neurons. If the coupling of neuromodulatory input to the SBC-like neurons was sufficiently strong, then these neurons could be activated by neuromodulatory inputs, even in the absence of thalamic inputs. We do not know whether this situation occurs *in vivo*.

*Omissions from the model.*

Many possibly relevant features were omitted from the model. These include: gap junction connections between eNGC neurons (*5*), activation of $GABA_B$ receptors, activation of muscarinic acetylcholine receptors (*6*), and possible feedback inhibition from deeper layer Martinotti cells (*7*). We did not consider a finer classification of L1 interneurons into sub-types. Our model also did not include cortico-cortical inputs. We did not study or simulate the effects of L1 interneuron activation on apical dendrites of deeper layer pyramidal cells. Activation of $5HT_3A$ ionotropic serotonin receptors is expected to have similar electrophysiological effects to activation of nicotinic acetylcholine receptors, though the distribution of these two receptor-types in the different L1 interneuron sub-classes may be different.

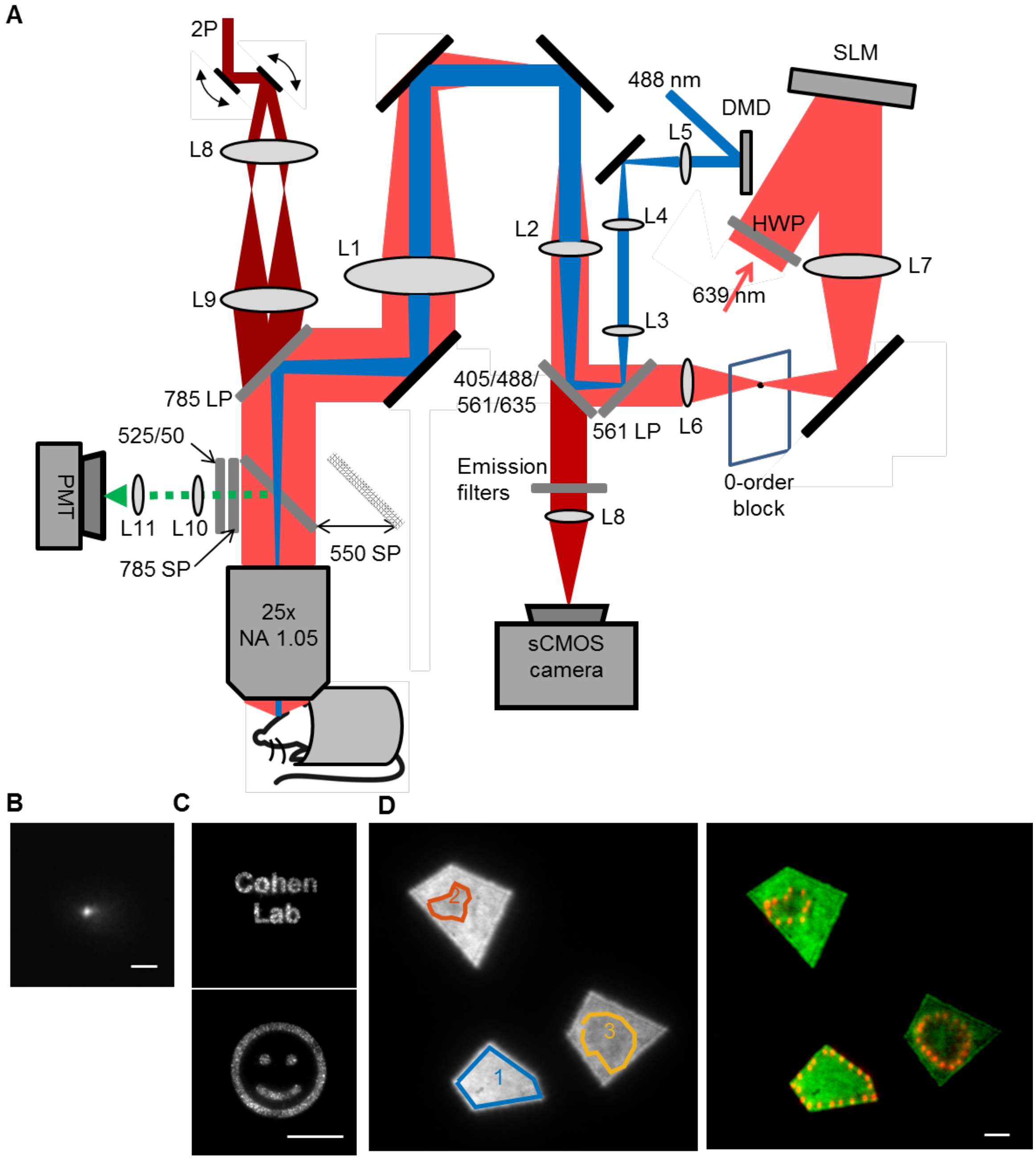


**Fig. S1. Instrument for all-optical electrophysiology *in vivo*.** (A) Layout of the optical setup. The spatial light modulator (SLM) diffractively patterned the red laser ($\lambda$ = 639 nm) into a set of discrete foci on the sample for holographic structured illumination excitation of a far red GEVI. The digital micromirror device (DMD) acted as a binary amplitude mask and projected a pattern of blue light onto the sample for targeted optogenetic activation of a blue-excited channelrhodopsin. Fluorescence from the sample was imaged onto a scientific CMOS camera. For two-photon (2P) imaging, pulsed infrared light ($\lambda$ = 920 nm) was scanned by a pair of galvo mirrors. A dichroic mirror was inserted into the beam path to direct green fluorescence onto a photomultiplier (PMT). Parts list in Table S3. Not shown: beam expansion and polarization control optics for each of the laser beams. (B) Point-spread function of the red illumination. Scale bar 10 μm. (C) Diffractively patterned red light illumination patterns projected onto a homogeneous fluorescent test sample. Scale bar 50 μm. (D) Combination of patterned blue and red illumination. Left: Patterns of fluorescence excited by blue light projected onto a homogeneous fluorescent test sample. Target patterns for the red illumination were manually defined. Right: Superposition of image of the red illumination on the green fluorescence excited by blue illumination. Scale bar 10 μm.

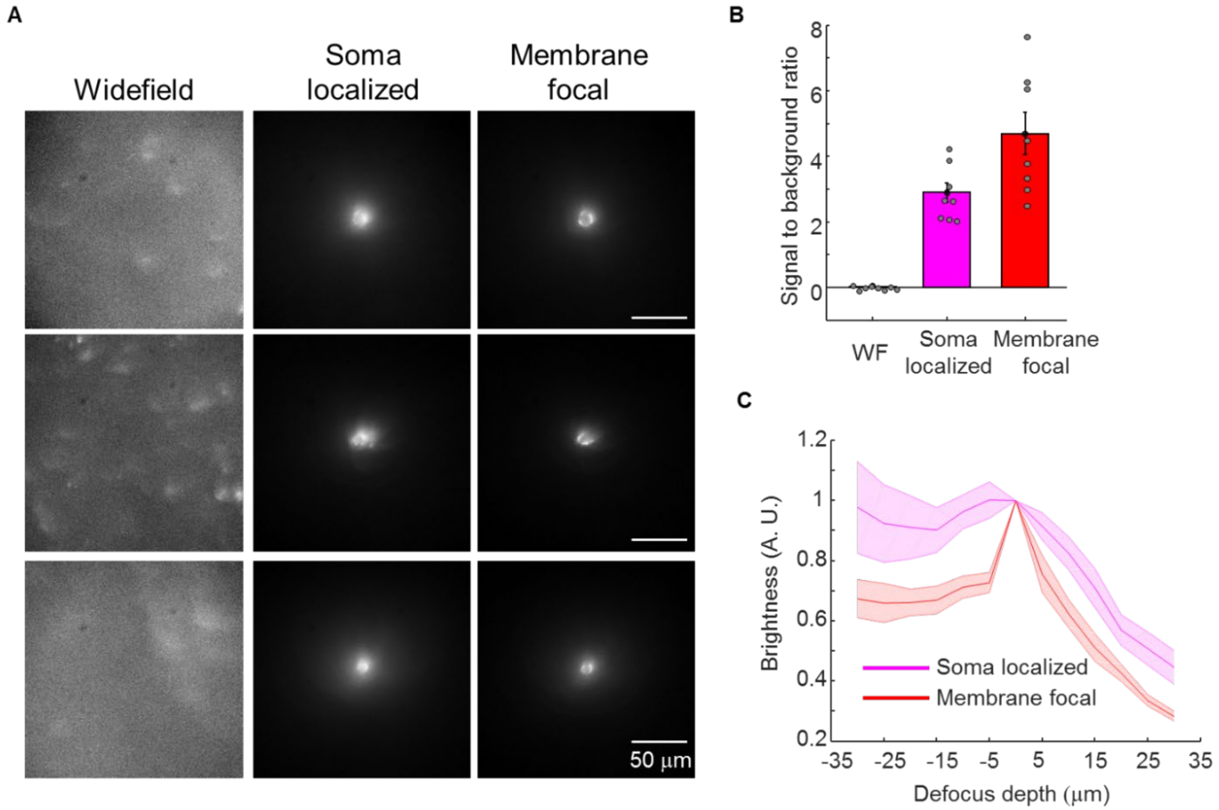


**Fig. S2. Holographic structured illumination microscopy improves signal-to-background ratio *in vivo*.** A) Representative images in the SomArchon fluorescence channel showing wide-field, cell-localized, and holographic focal illumination. B) Quantification of the signal (cell area) to background (surrounding region) ratio for the three illumination schemes. C) Quantification of the relative signal level as a function of defocus. Here a signal mask was defined on the in-focus image, and then applied to images taken at a series of defocus values.

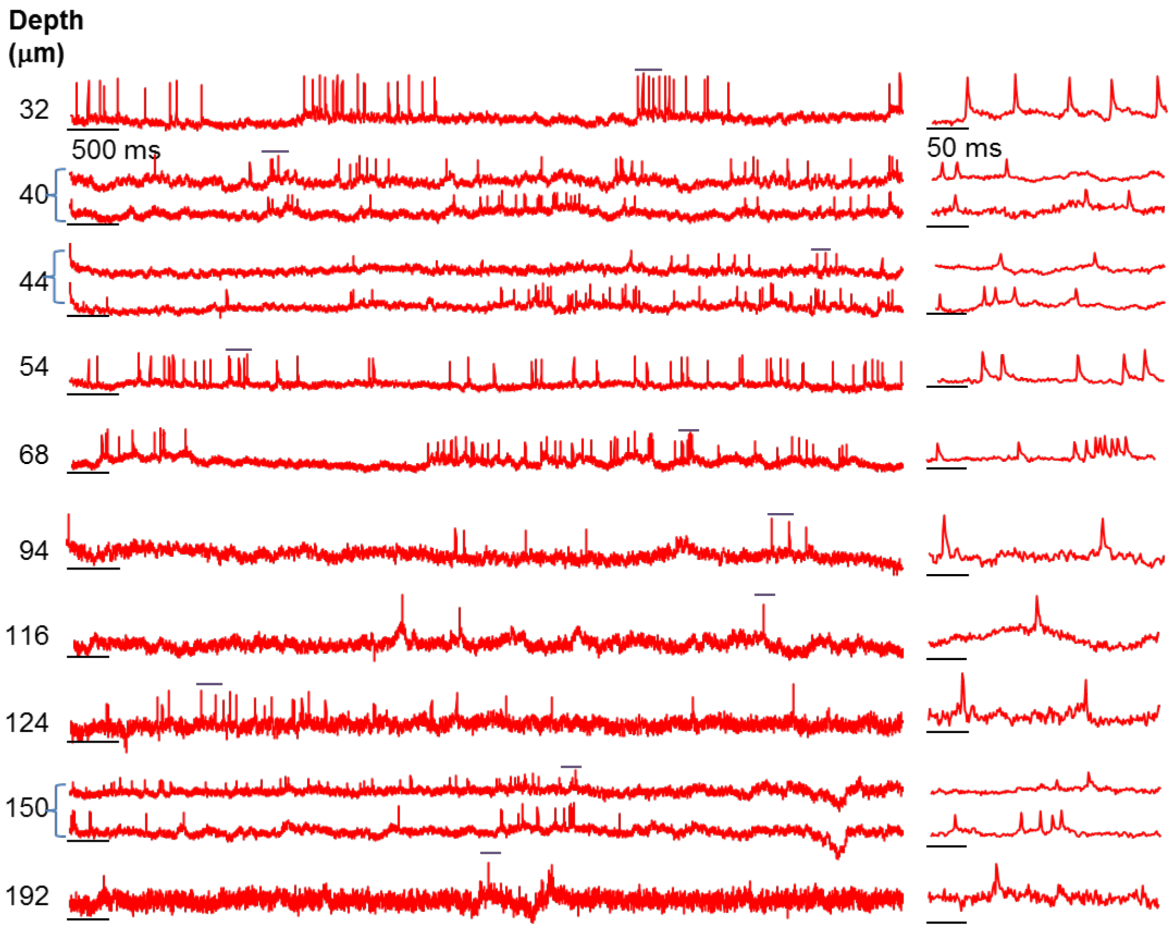


**Fig. S3. Voltage imaging in the cortical neurons at different depths in awake mice**. Fluorescence recordings of neurons expressing SomArchon-eGFP in barrel cortex of awake mice. Neurons were located via 2P microscopy and then targeted with holographic 1P red illumination (3 mW/cell). Fluorescence was recorded at 1 kHz. Left: single-trial recordings of spontaneous activity. Pairs of simultaneously recorded cells shown with brackets. Right: magnified views of the regions indicated by the purple over-bars from the left. SNR values (spike height:baseline noise) were 20 at the shallowest depth and 6.7 at the greatest depth.

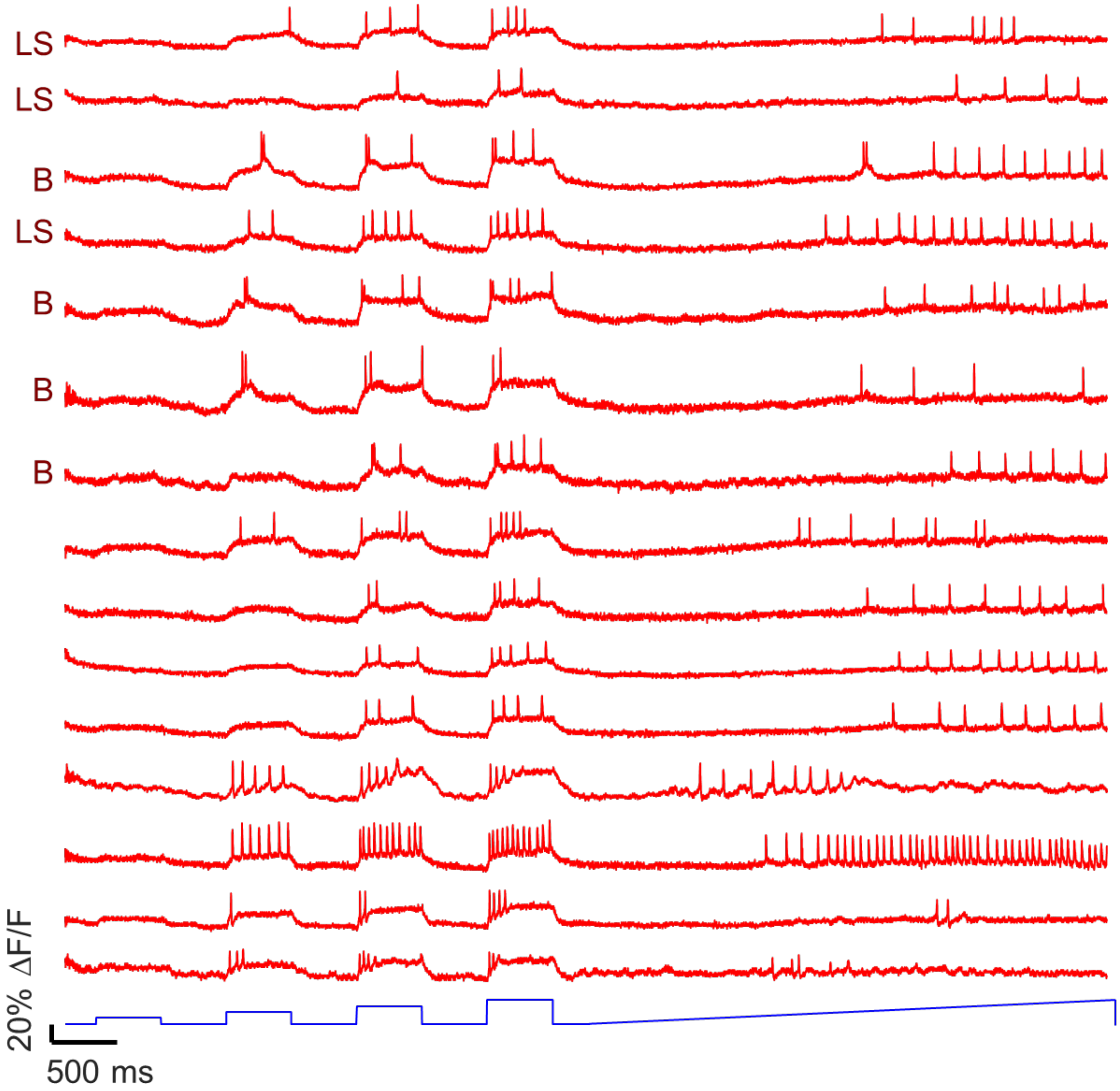


**Fig. S4. Optopatch excitability measurement of cortical L1 neurons in acute slices.** In cortical L1 neurons from a 5-HT3AR-Cre mouse expressing Optopatch4, the SomArchon fluorescence reported action potentials with high SNR. Red: fluorescence of SomArchon during optogenetic stimulation, recorded at 1 kHz. Blue: Optogenetic stimulus waveform (500 ms duration, 0.2 to 2.1 mW/mm$^2$, repeated at 1 Hz). The signal-to-noise ratio (spike:baseline noise) was 21 ± 1 in a 1 kHz bandwidth (n = 15 cells, 1 mW per cell, mean ± s.e.m.). Some of the spiking waveforms have been labeled as “Late spiking” (LS) or “Bursting” (B) in correspondence with established L1 firing phenotypes. Not all cells clearly fell into one of these classes.

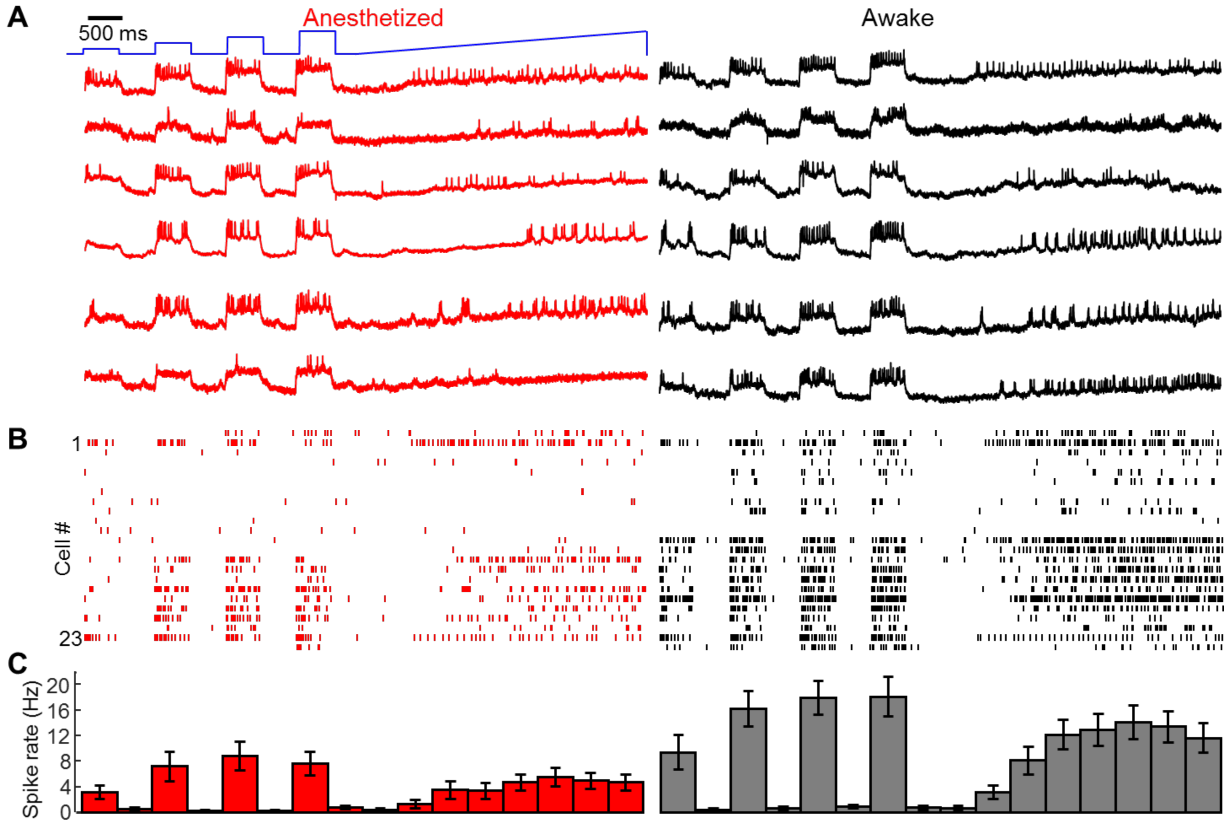


**Fig. S5. Paired recordings of L1 excitability under anesthesia and wakefulness.** (A) Individual L1 neurons were illuminated with steps of blue light (500 ms duration, 1.8 to 21 mW/mm$^2$, repeated at 1 Hz), followed by a ramp of blue light. Voltage was recorded at 1 kHz via holographic focused excitation of SomArchon fluorescence. Neuron coordinates were recorded relative to blood vessel landmarks. Anesthesia was then ended. After the animal awoke the measurements were repeated on the same set of cells. Each row represents repeated recordings of the same cell ($n$ = 23 neurons from 3 mice). (B) Spike raster for the complete dataset recorded under anesthesia and wakefulness. (C) Mean spike rate during each stimulation epoch. Error bars represent s.e.m..

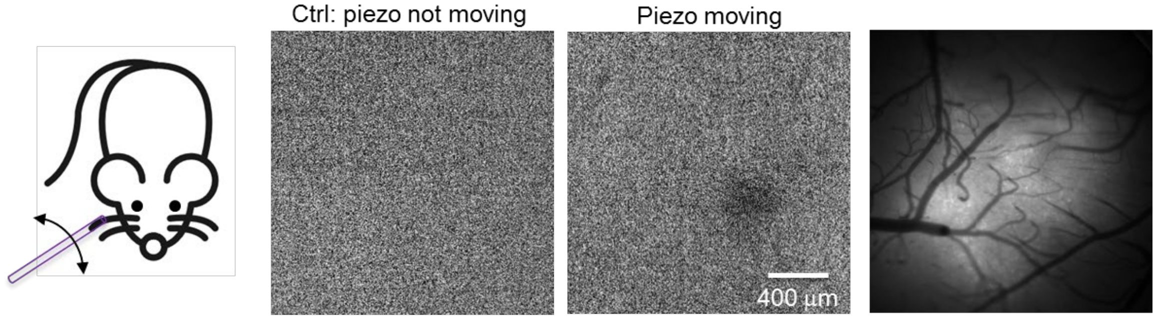


**Fig. S6. Intrinsic imaging to locate barrels corresponding to a single whisker.** In an anesthetized mouse, the surface of the brain was imaged via 640 nm reflected light. A single whisker was periodically stimulated (10 Hz, 4 s, followed by 16 s rest) for 5 min. Images were acquired at 10 Hz. The mean of the image acquired during the stimulated epochs was subtracted from the mean of the image acquired during the rest epochs. A dark spot highlighted the active barrel. A reference image taken with back-scattered blue light identified the blood vessel landmarks around the active barrel.

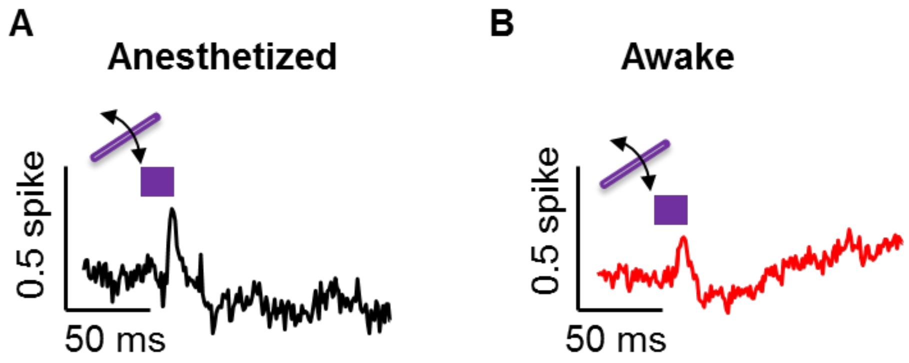


**Fig. S7. Whisker stimuli that failed to evoke spikes still evoked post-stimulus hyperpolarization.** Stimulus-triggered average waveforms were computed for stimulus events that failed to evoke a spike in the measured neuron. In both (A) anesthetized and (B) awake animals, the stimulus evoked an EPSP followed by an IPSP. Post-stimulus hyperpolarization in the absence of a spike indicates that the hyperpolarization arose from network inputs rather than from cell-autonomous mechanisms. Data from $n$ = 9 neurons (anesthetized) and $n$ = 16 neurons (awake), 3 mice in both cases.

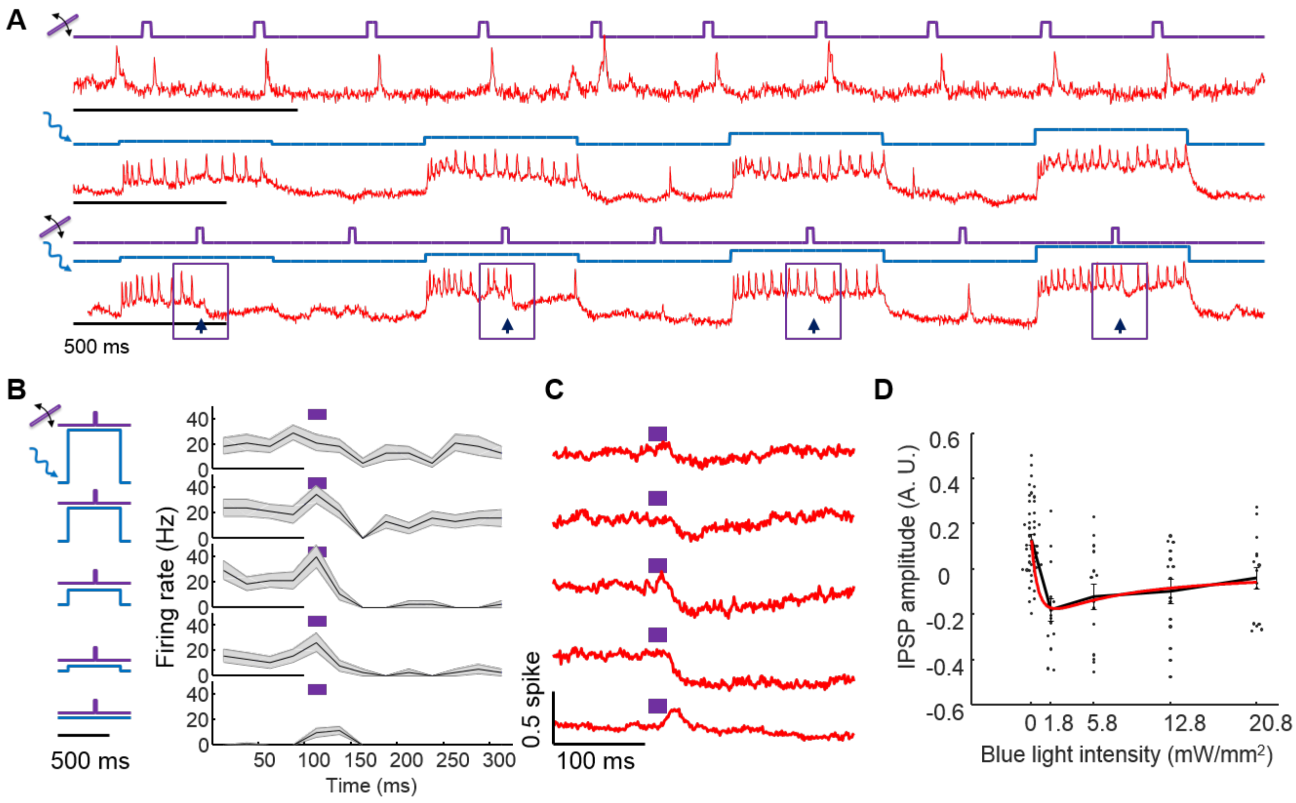


**Fig. S8. Optical dissection of E/I balance in L1 interneurons in anesthetized mice.** (A) Three recordings from a single neuron showing response to (top) whisker stimulus, (middle) optogenetic stimulus, and (bottom) simultaneous optogenetic and whisker stimuli. Arrows show whisker stimulus-evoked inhibition. (B) Mean spike rate evoked by whisker stimuli atop different levels of optogenetic stimulus. In the absence of optogenetic stimulation, whisker stimuli evoked single spikes. In the presence of optogenetic stimulation, whisker stimuli suppressed spiking. The suppression decreased in amplitude and duration as the strength of the optogenetic stimulus increased. Shading represents s.e.m. from $n$ = 15 neurons, 3 mice. (C) Mean whisker stimulus-evoked subthreshold waveforms at different levels of optogenetic drive. Spikes were digitally removed prior to averaging (Methods). (D) Comparison of IPSP amplitude as a function of optogenetic stimulus strength with numerical simulation from a simple conductance-based model.

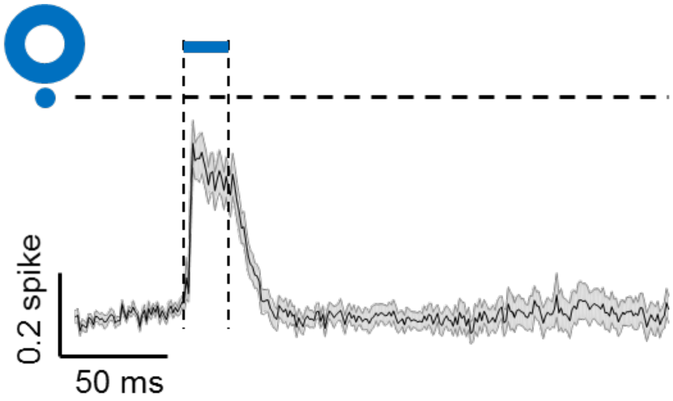


**Fig. S9. Light scatter contributes a depolarizing transient when a neuron is surrounded by a ring stimulus.** Mean fluorescence response of a central neuron during a 20 ms annular stimulus to surrounding neurons (25 mW/mm$^2$). This experiment is the same as in Fig. 3G, except that the central neuron is not subject to optogenetic stimulation. The experiment shown here and in Fig. 3G were performed on the same set of neurons in interleaved trials +/- central optogenetic depolarization (*n* = 25 neurons, 3 mice). The time-course of subthreshold depolarization matches the expectation from direct stimulation of the central neuron via scattered blue light from the surrounding annulus.

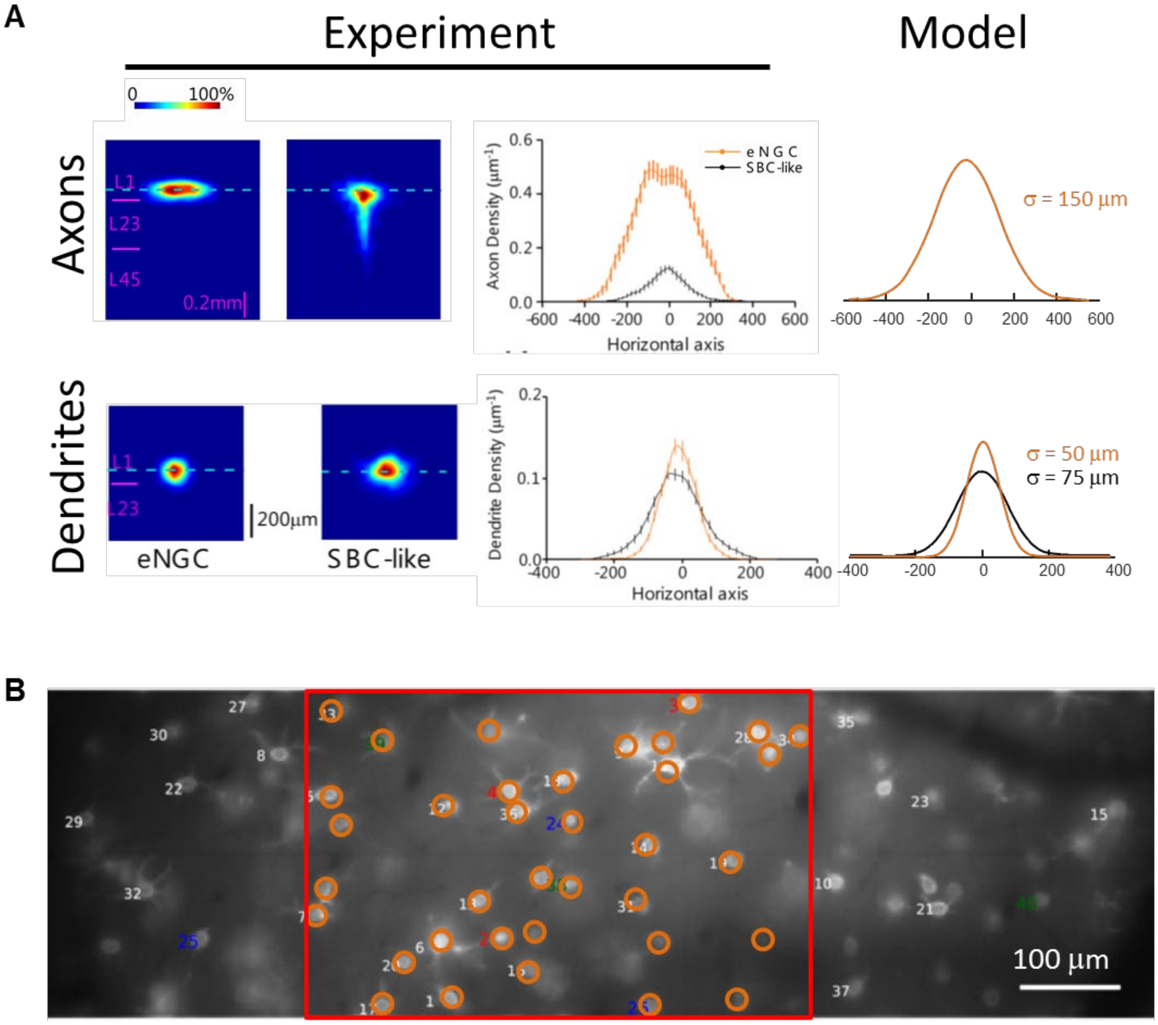


**Fig. S10. Calibration of morphological properties of L1 microcircuit.** A) Length-scale of connectivity. Data from (*8*) . Synaptic weights were set by gaussian distributions fit to the experimental data. The length-scale of each synaptic connection was set to the sum of the widths of the axonal and dendritic arbors. B) Density of neurons in L1. Cell density was estimated from published images of fluorescently labeled L1 interneurons. Data from(*9*) . These results were consistent with precise counts taken in the rat (*10*).

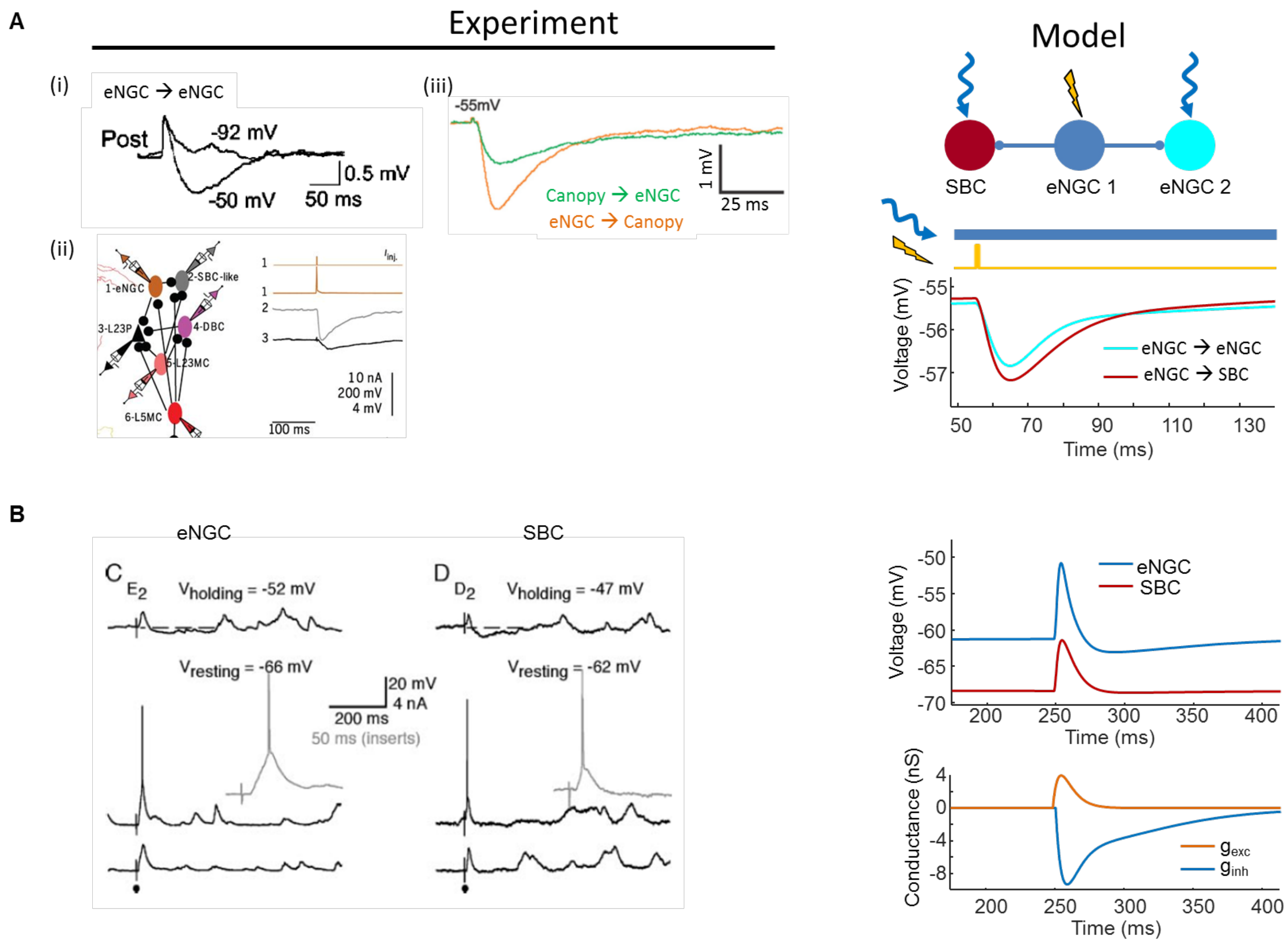


**Fig. S11. Calibration of synaptic properties of L1 model.** A) Calibration of inhibitory synaptic strengths. Data from (i) (*5*) (ii) (*8*) (iii) (*11*) . B) Model for synaptic inhibition. An eNGC cell was stimulated to spike once. The timecourse and amplitude of the postsynaptic inhibitory conductances were adjusted to match approximately the experimental IPSPs. In the simulation, the postsynaptic cells were optogenetically depolarized to -55 mV to introduce a driving force for $Cl^-$ entry. B) Calibration of excitatory thalamocortical synaptic strengths. Patch clamp recordings showed that whisker stimuli evoked EPSPs of 3 – 7 mV. In the model, excitatory synaptic strength was adjusted so that, in combination with sensory-induced lateral inhibition, the EPSP from baseline approximately matched the timecourse and amplitude observed experimentally.

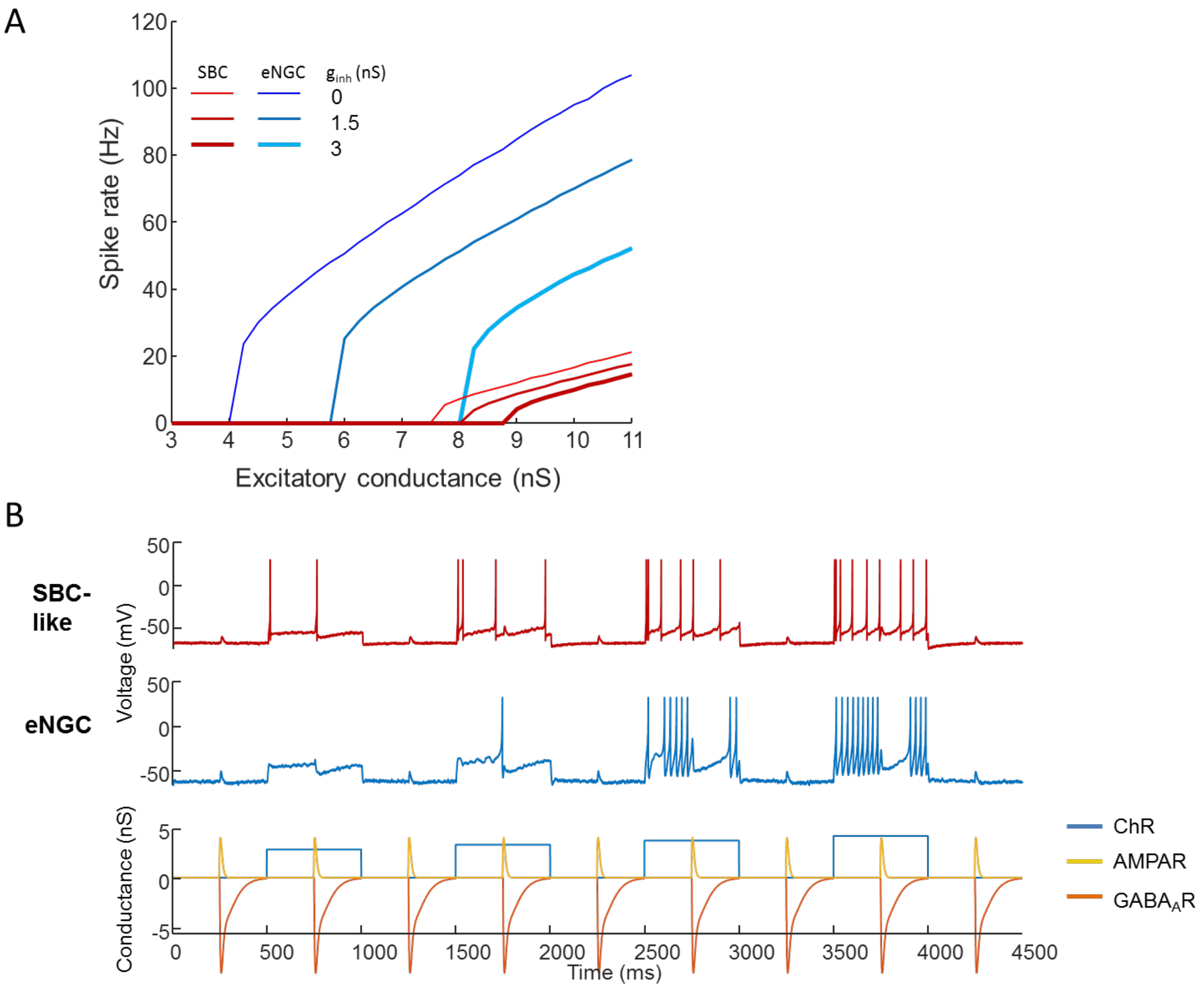


**Fig. S12. Characterization of single-cell firing properties in L1 model.** A) Steady-state spiking rate as a function of excitatory and inhibitory conductance. eNGC cells showed a clear threshold to activate spiking, while SBC cells showed a more graded response. The difference in activation threshold implied that the SBC output cells were never activated without simultaneous activation of the eNGC cells, i.e. that excitation of the output always occurred in the presence of lateral inhibition. $VIP^+$ interneurons (associated with the SBC-like population) have been shown experimentally to have a more hyperpolarized resting potential and a more depolarized threshold potential than other L1 interneuron classes (*11*) . B) Numerical simulations in which single cells were exposed to step-wise increases in channelrhodopsin activation and paired excitatory and inhibitory inputs. In the absence of optogenetic stimulation, the synaptic inputs drove small depolarization. In the presence of optogenetic stimulation, the synaptic inputs drove primarily inhibition. The magnitude and duration of the inhibition decreased as the strength of the optogenetic stimulation increased.

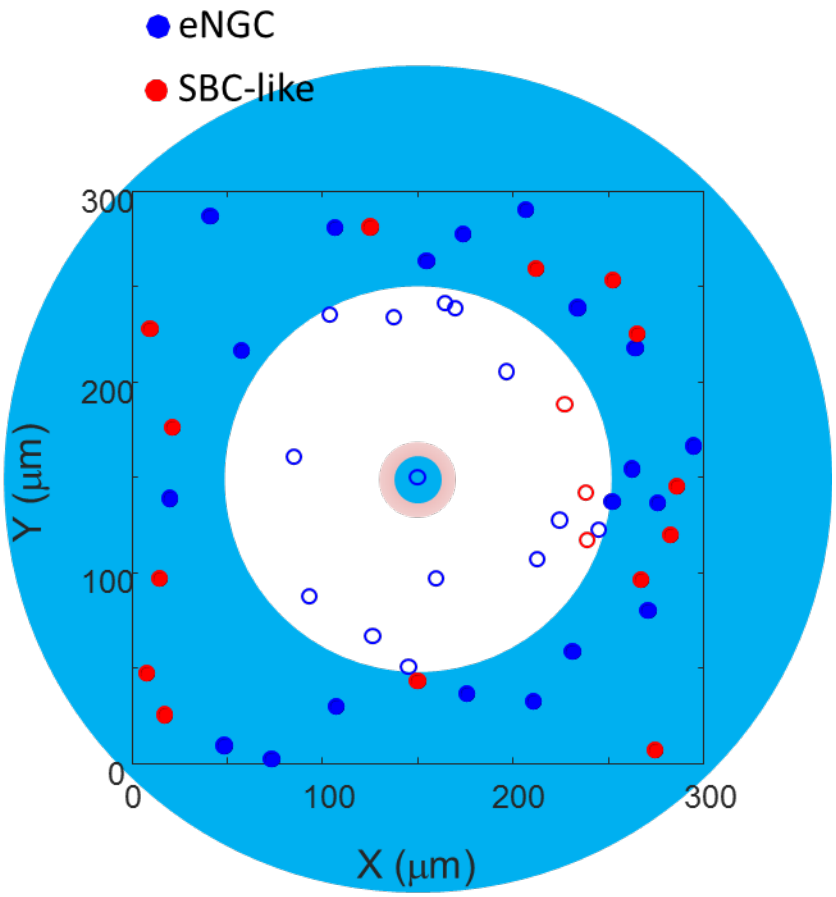


**Fig. S13. Geometry of annular optogenetic stimulation in a simulated L1 barrel.** Cells were randomly positioned in a region 300 µm on an edge, 150 µm deep. A single cell (eNGC or SBC-like) was manually defined to reside at the center of the region. The central neuron was subjected to tonic optogenetic depolarization. Neurons at radii $r > 100$ µm from the center were subjected to pulsed optogenetic stimulation. The voltage in the central neuron was plotted in Fig. 5D.

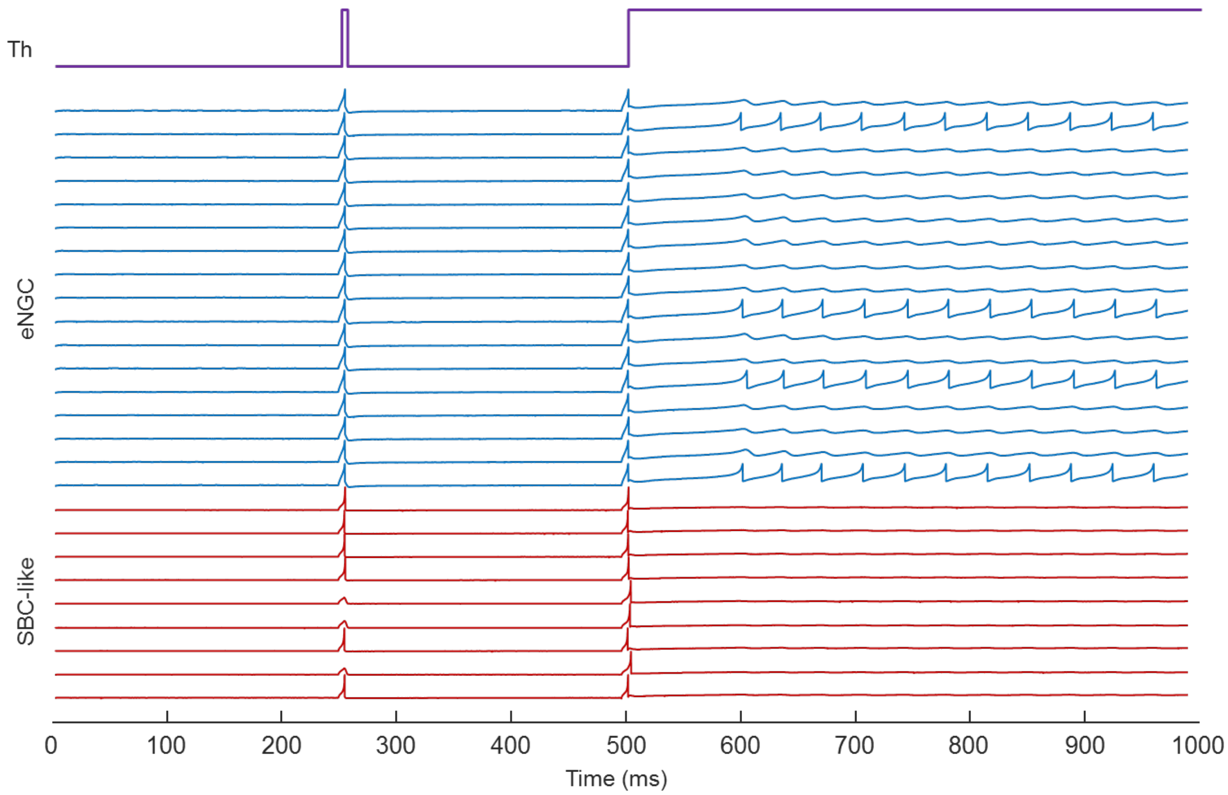


**Fig. S14. Comparison of L1 network responses to abrupt vs. sustained inputs.** A thalamic input was delivered simultaneously to all eNGC and SBC-like neurons in an L1 simulation. Transient inputs activated all eNGC neurons and most SBC-like neurons. Sustained inputs of the same strength initially activated most neurons. After a period of global network inhibition, a subset of eNGC neurons became tonically active, suppressing activation of the other eNGC neurons and of the SBC-like population. Here a randomly selected subset of the neurons are plotted.

| Parameter | eNGC neuron | SBC-like neuron | Meaning and comments |
|---|---|---|---|
| C | 30 pF | 30 pF | Membrane capacitance. Artificially low compared to patch clamp recordings, necessary for fast spikes. |
| k | 0.3 pA/mV | 1.2 pA/mV | Conductance of endogenous voltage-gated channels (pA/mV = nS) |
| $V_r$ | -66 mV | -70 mV | Resting potential |
| $V_t$ | -36 mV | -39 mV | Threshold potential |
| $V_p$ | 30 mV | 30 mV | Spike peak potential |
| a | 0.17 $ms^{-1}$ | 0.01 $ms^{-1}$ | Recovery rate constant |
| b | 5 pA/mV | 6 pA/mV | Recovery conductance |
| c | -45 mV | -65 mV | Post-spike reset value for membrane voltage |
| d | 100 pA | 90 pA | Post-spike jump in adaptation variable, *u*. |
| $G_{ds}$ | 1.2 pA/mV | - | Dendrite-to-soma coupling strength |
| $G_{sd}$ | 0.01 pA/mV | - | Soma-to-dendrite coupling strength |
| $V_{ChR}$, $V_{AMPA}$, $V_{nAchR}$ | 0 mV | 0 mV | Reversal potential of excitatory conductances |
| $V_{Cl}$ | -70 mV | -70 mV | Reversal potential of inhibitory ($GABA_A$R-mediated) conductances |

**Table S1. Parameters of Izhikevich-type models of L1 interneurons.** The eNGC cells were described by a two-compartment cell, with one active compartment and a passive dendrite. The SBC-like cells were described by a one-component model. Parameters were adjusted by hand to match the observed firing patterns.

| Parameter | Source Data | Model |
|---|---|---|
| eNGC firing pattern | Chu et al. J. Neurosci. 23 (2003): 96-102. Fig. 1 | Modified from Izhikevich, Dynamical Systems in Neuroscience, Eq. 8.28 |
| SBC firing pattern | Chu et al. J. Neurosci. 23 (2003): 96-102. Fig. 1 | Modified from https://www.izhikevich.org/publications/figure1.m Fig. 1E,F |
| IPSP waveform: eNGC → eNGC | Chu et al. J. Neurosci. 23 (2003): 96-102. Fig. 5 | $g_{inh}(t) = g_{inh}^{0}\left(\frac{t}{\tau_1}e^{1-\frac{t}{\tau_1}} + 0.6\frac{t}{\tau_2}e^{1-\frac{t}{\tau_2}}\right)$ <br> $g_{inh}^{0}$ = 1 nS, $t_1$ = 7 ms, $t_2$ = 35 ms |
| IPSP amplitude eNGC → eNGC | Jiang et al. Science 350 (2015): aac9462. Table S6 | -1.6 mV |
| IPSP waveform eNGC → SBC | Chu et al. J. Neurosci. 23 (2003): 96-102. Fig. 4 Jiang et al. Science 350 (2015): aac9462. Fig. 3a | $g_{inh}(t) = g_{inh}^{0}\left(\frac{t}{\tau_1}e^{1-\frac{t}{\tau_1}} + 0.6\frac{t}{\tau_2}e^{1-\frac{t}{\tau_2}}\right)$ <br> $g_{inh}^{0}$ = 1.6 nS, $t_1$ = 7 ms, $t_2$ = 35 ms |
| IPSP amplitude eNGC → SBC | Jiang et al. Science 350 (2015): aac9462. Table S6 | -1.8 mV |
| Length scale of eNGC → eNGC coupling | Jiang et al. Science 350 (2015): aac9462. Figs. S2, S3 | s = 200 µm (sum of axonal and dendritic length scales) |
| Length scale of eNGC → SBC coupling | Jiang et al. Science 350 (2015): aac9462. Figs. S2, S3 | s = 225 µm (sum of eNGC axonal and SBC dendritic length scales) |
| L1 neuron density | Abdelfattah et al., BioRxiv: /10.1101/436840. Fig. S23 Meyer et al., PNAS 110 (2013) 19113-19118. Table S2 | 3000-7,000 $mm^{-3}$, 15 – 67 neurons/barrel (in rat) |
| Ratio of eNGC to SBC cells | Schuman et al. J. Neurosci. 39 (2019): 125-139. Fig. 2,4 | ~2:1 |
| Thalamocortical EPSP waveform in eNGC | Zhu and Zhu, J. Neurosci. 24 (2004): 1272-1279. Fig. 2 | $g_{exc}(t) = g_{exc}^{0}\frac{t}{\tau}e^{1-\frac{t}{\tau}}$ <br> $g_{exc}^{0}$ = 4 nS, t = 6 ms |
| Thalamocortical EPSP amplitude in eNGC | Zhu and Zhu, J. Neurosci. 24 (2004): 1272-1279. Fig. 2 Lee et al., J. Neurosci. 30 (2010): 16796-16808. Fig. 7 | 3 - 7 mV |
| Thalamocortical EPSP waveform in SBC | Zhu and Zhu, J. Neurosci. 24 (2004): 1272-1279. Fig. 2 | $g_{exc}(t) = g_{exc}^{0}\frac{t}{\tau}e^{1-\frac{t}{\tau}}$ <br> $g_{exc}^{0}$ = 4 nS, t = 6 ms |

| | | |
|---|---|---|
| Thalamocortical EPSP amplitude in SBC | Zhu and Zhu, J. Neurosci. 24 (2004): 1272-1279. Fig. 2<br>Lee et al., J. Neurosci. 30 (2010): 16796-16808. Fig. 7<br>(But see also Cruikshank et al. J. Neurosci. 32 (2012): 127813-17823. Fig. 4d) | 3 - 7 mV |
| Strength of neuromodulatory input to eNGC | Unknown | Assumed to be same for eNGC and SBC |
| Strength of neuromodulatory input to SBC | Unknown | Assumed to be same for eNGC and SBC |

**Table S2.** Parameters used in simulations of L1 network activity. Some parameters were from data in rats or from different brain regions. We verified that where there was uncertainty in parameter values, the simulation results did not qualitatively depend on precise parameter values.

| Item | Part number | Comments |
|---|---|---|
| 2P Laser | Coherent DeepSee | |
| Scanning galvos | Cambridge Technologies 6215H | |
| Scan lens | Thorlabs, SL50-CLS2 | |
| 2P tube lens | Thorlabs, TL200-CLS2 | |
| PMT | Hamamatsu, H11706P-40 | |
| 25x objective | Olympus XLPLN25XWMP2 | |
| L1 | 400 mm, Edmund, 88-598-INK | |
| L2 | Effective focal length, 150 mm, two 300 mm lenses, Thorlabs, AC508-300-A-ML | Two lenses back to back in Plössl configuration to reduce aberration |
| L3 | 75 mm, Thorlabs, AC254-075-A-ML | Relay lens to allow more space |
| L4 | 75 mm, Thorlabs, AC254-075-A-ML | Relay lens to allow more space |
| L5 | 60 mm, Thorlabs, AC254-060-A-ML | |
| L6 | First generation: Sigma macro 18-200 mm<br><br>Second generation: 75 mm, Thorlabs, AC508-075-A-ML | First generation: Zoom lens for varying NA of red illumination and size of red targeted region;<br><br>Second generation: 75 mm lens to reduce aberration |
| L7 | 200 mm, Thorlabs, AC508-200-A-ML | |
| L8 | 45 mm, Olympus XLFLUOR 4X/340 | Objective used as a tube lens for reducing aberration and achieving large field of view |

| | | |
|---|---|---|
| L9 | Thorlabs, SL50-CLS2 | Optimized scan lens |
| L10 | Thorlabs, TL200-CLS2 | Optimized tube lens |
| L11 | 16 mm, Thorlabs, AC080-016-A-ML | |
| L12 | 75 mm singlet, Thorlabs, LA1608-A-ML | |
| 488 nm laser | Cobolt, 06-01 series, $\lambda$ = 488 nm, 60 mW | |
| AOTF | Gooch and Housego TF525-250-6-3-GH18A | |
| DMD | Vialux, V-7001 VIS | |
| 639 nm laser | CNI Inc., MRL-FN-639, $\lambda$ = 639 nm, 700 mW single transverse mode | |
| SLM | Meadowlark 1920SLM VIS | |
| 0-order block | home-made anti-pinhole comprised of a dot of solder on a glass slide | |
| sCMOS camera | Hamamatsu ORCA-Flash 4.0 | |
| Sample stage | Sutter instrument, FG-MPC78, Moving stage plat W/MPC-200 for XY stage; SA-MP285-1X-M for Z axis | |
| DAQ system | NI PCIe-6363 | |

**Table S3.** Components required to build an optical system for holographic structured illumination voltage imaging combined with patterned optogenetic stimulation.
